# DDHD2 interacts with STXBP1 to mediate long-term memory via the generation of myristic acid

**DOI:** 10.1101/2023.05.11.540316

**Authors:** Isaac O. Akefe, Benjamin Matthews, Saber H. Saber, Bharat G. Venkatesh, Rachel S. Gormal, Daniel G. Blackmore, Emma Sieriecki, Yann Gambin, Jesus Bertran-Gonzalez, Alysee A. Michaels, Mingshan Xue, Benjamin Cravatt, Merja Joensuu, Tristan P. Wallis, Frédéric A. Meunier

**Affiliations:** Clem Jones Centre for Ageing Dementia Research, Queensland Brain Institute, The University of Queensland, St Lucia, QLD, 4072, Australia; School of Medical Science, University of New South Wales, Randwick, NSW, 2052, Australia; EMBL Australia, Single Molecule Node, University of New South Wales, Sydney, 2052, Australia; Decision Neuroscience Lab, School of Psychology, UNSW Sydney, Australia; Department of Neuroscience, Baylor College of Medicine, Houston, TX, United States; The Cain Foundation Laboratories, Jan and Dan Duncan Neurological Research Institute at Texas Children’s Hospital, Houston, TX, United States; Department of Molecular and Human Genetics, Baylor College of Medicine, Houston, TX, United States; The Department of Chemistry and The Skaggs Institute for Chemical Biology, The Scripps Research Institute, La Jolla, CA, United States; The School of Biomedical Sciences, The University of Queensland, St Lucia, QLD, 4072, Australia

**Keywords:** Lipids, Phospholipase A1, Free Fatty Acids, Myristic Acid, Learning and Memory

## Abstract

The phospholipid and free fatty acid (FFA) composition of neuronal membranes plays a crucial role in learning and memory, but the mechanisms through which neuronal activity affects the brain’s lipid landscape remain largely unexplored. Saturated FFAs, particularly myristic acid (C14:0), strongly increase during neuronal stimulation and memory acquisition, suggesting the involvement of phospholipase A1 (PLA1) activity in synaptic plasticity. Here, we show that genetic ablation of the DDHD2 isoform of PLA1 in mice reduced memory performance in reward-based learning and spatial memory models prior to the development of neuromuscular deficits, and markedly reduced saturated FFAs across the brain. DDHD2 was shown to bind to the key synaptic protein STXBP1. Using STXBP1/2 knockout neurosecretory cells and a haploinsufficient *STXBP1*^+/-^ mouse model of STXBP1 encephalopathy that is also associated with intellectual disability and motor dysfunction, we show that STXBP1 controls the targeting of DDHD2 to the plasma membrane and the generation of saturated FFAs in the brain. Our findings suggest key roles for DDHD2 and STXBP1 in the lipid metabolism underlying synaptic plasticity, learning and memory.

## Introduction

Activity-dependent changes to the brain’s lipid landscape are widely believed to play a role in neuronal activity, learning and memory. Uncovering the mechanism by which molecular and cellular interactions between lipids and specific proteins at the synapse contribute to memory formation and consolidation remains crucial in establishing a deeper understanding of synaptic plasticity and bases for neurological diseases (Wenk & De Camilli, 2004). In addition to acting as building blocks of synaptic vesicles and neuronal plasma membranes, brain lipids are involved in diverse neuronal functions including scaffolding (Lee *et al*, 2020), neurodevelopment (Zhu *et al*, 2016), signalling (Falomir-Lockhart *et al*, 2019), neuroinflammation (Giacobbe *et al*, 2020), neurometabolism (Bruce *et al*, 2017), and cognition (de Mendoza & Pilon, 2019; Derbyshire, 2018; Egawa *et al*, 2016; Joffre, 2019; Snowden *et al*, 2017; Tracey *et al*, 2018). Dynamic regulation of the brain-lipid landscape via the coordinated activity of phospholipases and other enzymes is crucial to mediate appropriate membrane curvature, surface chemistry, fluidity and fusogenicity of the phospholipid membrane bilayer during vesicular trafficking, exocytosis, endocytosis and synaptic plasticity underpinning memory formation (Alrayes *et al*, 2015; Barber & Raben, 2019; Bruce *et al*., 2017; Carta *et al*, 2014; Zhu *et al*., 2016). The importance of phospholipases in this process is exemplified in hereditary spastic paraplegia (HSP; Strümpell-Lorrain disease) in which a mutation in the *DDHD2* gene of phospholipase A1 (PLA1) is associated with neuromuscular and cognitive dysfunction (Alrayes *et al*., 2015; Bertran-Gonzalez *et al*, 2013; Inloes *et al*, 2014; Joensuu *et al*, 2020; Murala *et al*, 2021), autism (Matoba *et al*, 2020), schizophrenia (Park *et al*, 2021), intellectual disability (Alrayes *et al*., 2015), and mental retardation (Alrayes *et al*., 2015; Bertran-Gonzalez *et al*., 2013; Darios *et al*, 2020; Inloes *et al*., 2014; Joensuu *et al*., 2020). Although this establishes a potential causal link between PLA1 and cognitive function, the mechanisms by which phospholipases affect the lipid landscape to mediate synaptic plasticity that is conducive to memory acquisition in the brain are unknown.

Phospholipids and their free fatty acid (FFA) metabolites (Kihara *et al*, 2014) have been shown to bind key exocytic proteins such as syntaxin-1A (STX1A), and Munc18-1 (STXBP1) (Jang *et al*, 2012; Lee *et al*, 2004), to regulate the rate of synaptic vesicle exo- and endocytosis. Phospholipids and FFAs also function alone, as signalling molecules, or as reservoirs of lipid messengers which, in turn, regulate and interact with several signalling cascades, contributing to synaptic structure and function (Montaner *et al*, 2018; Paoletti *et al*, 2016; Wang *et al*, 2016). Polyunsaturated fatty acids (PUFAs) which contain more than one double bond in their backbone acyl tail, such as arachidonic acid (AA; C20:4) and docosahexaenoic acid (DHA; C22:6), play a vital role in soluble N-ethylmaleimide-sensitive factor attachment protein receptor (SNARE)-mediated synaptic transmission (Sidhu *et al*, 2016). Specifically, the non-covalent interaction of AA has been shown to allow STX1A to form the SNARE complex (Connell *et al*, 2007). Furthermore, FFAs are capable of regulating neurotransmission by modulating membrane curvature, fluidity, and fusogenicity, consequently making the membrane architecture more amenable to vesicle fusion (Graham & Kozlov, 2010). Cholesterol, diacylglycerol (DAG), and phosphatidic acid (PA) are cone-shaped lipids which can also induce negative membrane curvature to promote membrane fusion (Ammar *et al*, 2013).

During neurological processes such as neurotransmitter release, long-term potentiation and subsequent synaptic plasticity, the targeted activity of phospholipases plays a key role by catalysing the hydrolysis of phospholipids to generate FFAs thereby changing the local environment of the targeted bilayer (Joensuu *et al*., 2020; Puchkov & Haucke, 2013) and consequently impacting neurotransmission (Goldschmidt *et al*, 2016). Phospholipase A2 (PLA2) hydrolyses the fatty acyl chain on the *sn-2* position of canonical phospholipids to generate unsaturated FFAs such as AA, which further initiate a cascade of bioactive signalling molecules in many biological systems including neuroinflammation, neurotransmitter release and long-term potentiation (Higgs & Glomset, 1996; Inloes *et al*., 2014).

With the advent of unbiased lipidomic approaches, the involvement of PLA1 family members was suggested as secretagogue stimulation of neuroexocytosis was primarily shown to trigger a major increase in saturated FFAs in cultured neurons and neurosecretory cells (Narayana *et al*, 2015). Further, acquisition/consolidation of long-term fear memory in rats also correlates with strong increases in saturated FFAs (Wallis *et al*, 2021). These results pointed to a role in memory acquisition of PLA1 family members via the production of saturated FFAs and dynamic alteration of the phospholipid landscape in the brain. The importance of PLA1 to neuronal function is further substantiated in genetic disorders in which a prominent isoform of PLA1, called DDHD2, is absent or mutated. While DDHD2 is highly expressed in the central nervous system, the understanding of its role in neuronal function and plasticity is limited (Alrayes *et al*., 2015; Inoue *et al*, 2012; Joensuu *et al*., 2020).

To investigate the function of DDHD2 in mediating synaptic plasticity and memory formation, we used a DDHD2 knockout mouse model of hereditary spastic paraplegia (HSP) (Inloes *et al*., 2014). We employed a number of neurobehavioural paradigms to track the onset and progression of both neuromotor and cognitive decline throughout the lifespan of the *DDHD2^-/-^* mice. These longitudinal experiments were conducted on 3mo young mice that were largely asymptomatic, and again at 12mo, when the animals exhibit typical symptoms of neuromuscular dysfunction. We assessed the FFA changes in response to an instrumental conditioning paradigm that depends on reward-based associative memory. We useda rapid isotope-based multiplexing FFA analysis pipeline (FFAST (Wallis *et al*., 2021)) to quantify, in conditioned animals, the responses of 19 FFAs across 5 brain regions involved with cognitive and neuromotor functions. We found that reward-based instrumental conditioning can drive brain region-specific changes in the brain lipidome, characterised by increases in saturated FFAs, analogous to the response reported for a fear-based memory paradigm (Wallis *et al*., 2021). In contrast, *DDHD2* knockout mice exhibited a significant reduction in the response of saturated FFAs (especially myristic acid) even prior to the onset of memory impairment. This suggests that saturated FFA responses driven by DDHD2 may be a feature of memory acquisition in general, and are likely coupled to the activity of proteins involved in synaptic function. We demonstrate that DDHD2 primarily interacts with STXBP1 (also referred to as Munc18-1), a key pre-synaptic exocytic protein with chaperone functions (Arunachalam *et al*, 2008; Han *et al*, 2011). We reveal that this interaction controls the transport of DDHD2 to the plasma membrane and the subsequent generation of saturated FFAs. The role of STXBP1 in neuronal FFA metabolism was further demonstrated by the reduction of FFA levels in STXBP1 knockout neurosecretory cell lines and across the brains of heterozygous STXBP1 knockout mice which is a model of STXBP1 encephalopathy (Chen *et al*, 2020). Together these data suggest a novel role for the *DDHD2* isozyme of PLA1 in regulating FFA changes associated with memory, via interaction with STXBP1.

## Results

### Instrumental conditioning drives region-specific changes in the brain lipidome

The predominant increase in saturated FFA that occurred during fear conditioning suggested that this response was associated with the process of memory formation in the brain (Wallis *et al*., 2021). To assess whether this was a general response to learning, we tested whether it was associated with other memory paradigms. Instrumental conditioning is a long-term learning process that depends on associative memories linking a behavioral response with reward (Fig. 1A), and is known to initiate molecular and cellular mechanisms liable for memory formation in the posterior striatum and prefrontal cortex, amongst other regions (Balleine, 2019). Over the entire 17 days of training, the instrumental animals learned to push a lever for a food reward, while the control animals received no reward regardless of lever pushing. This was reflected by a considerably greater number of lever presses (P < 0.001) in the instrumental mice compared to the control cohort which exhibited essentially no lever presses (Fig. 1B, Extended Data Fig. 1A).

**Fig. 1.**
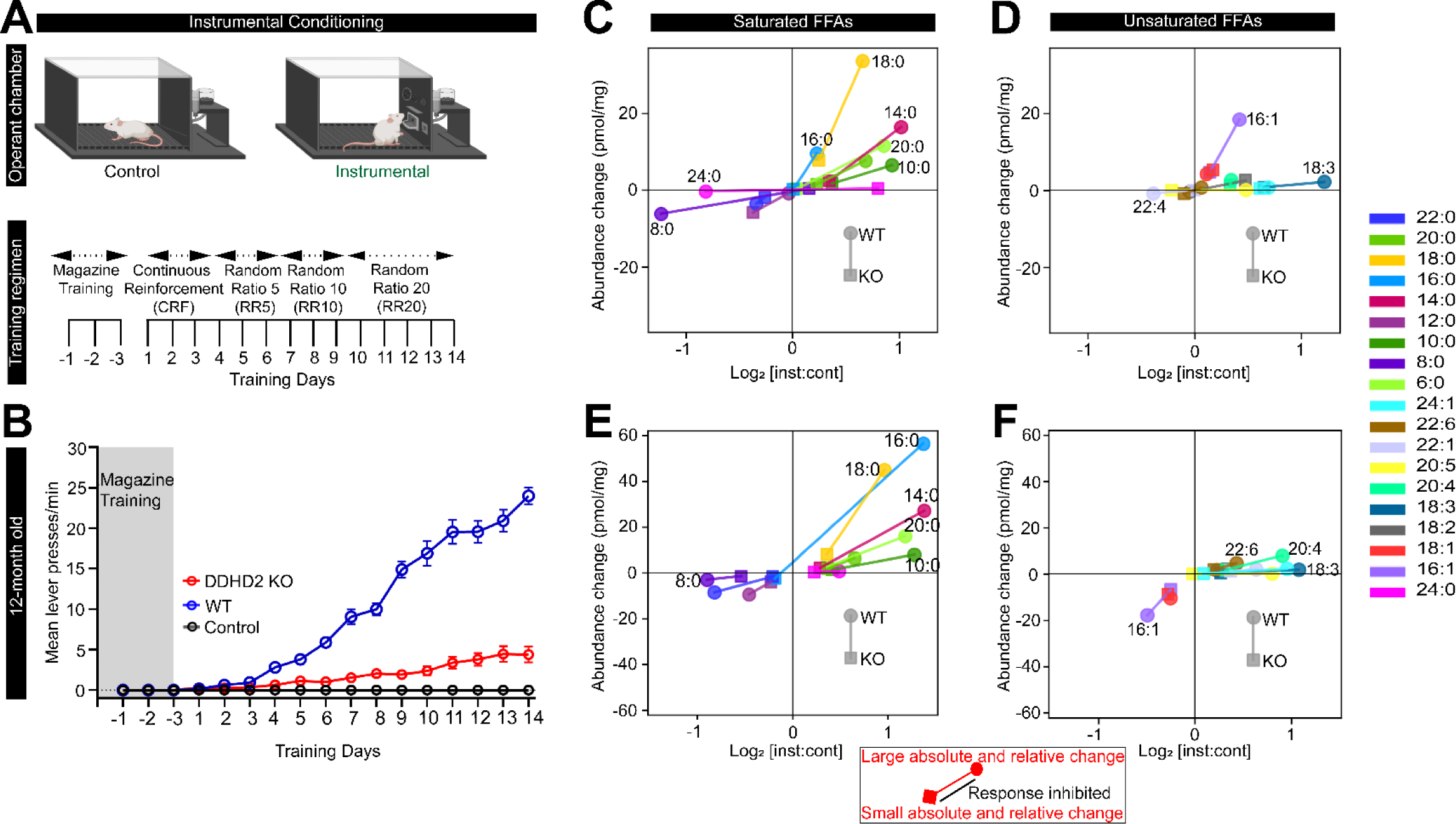
Behavioural and FFA responses to instrumental conditioning in *DDHD2^+/+^* vs *DDHD2^-/-^*mice. Instrumental conditioning was performed in cohorts of *DDHD2^+/+^* (WT) or *DDHD2^-/-^* (KO) mice vs controls respectively. **A** Illustration of the procedure for operant lever conditioning in instrumental vs control mice; n=20 (figure was produced using biorender). **B** Line graph showing mean lever presses per minute in 12mo WT vs KO mice (n=20 each; for instrumental and control cohorts of animals). Following conditioning animals were euthanised and brains were collected for FFAST analysis. Linked scatterplots show the absolute change (y axis) and log_2_ fold change (x axis) in mean concentration for each FFA in WT (round symbol) and KO (square symbol) mice. **C** Saturated FFA and, **D** Unsaturated FFA responses to instrumental conditioning within the dorsal hippocampus of 3mo WT vs KO mouse brains (n=6). **E** Saturated FFA and, **f** Unsaturated FFA responses to instrumental conditioning in the dorsal hippocampus of 12mo WT vs KO mouse brains (n=6).

FFAs from each condition were solvent extracted from homogenised tissue and independently derivatized using FFAST, and combined along with labelled FFA standards prior to multiplex tandem liquid chromatography mass spectrometry (LC-MS/MS). FFAST fragment peak intensities relative to the internal standard were used to determine absolute abundance of the 19 FFAs in each condition (Extended Data Fig. 1D; Table S1, S2, and S3).

Consistent with the preserved intrinsic architecture of the 5 brain regions examined, we uncovered considerable heterogeneity in the concentration of FFAs across the assayed brain regions, with the highest concentrations relative to tissue weight in the prefrontal cortex, and lowest concentrations in the cerebellum. Following instrumental conditioning, we uncovered an overall increase in the FFA levels across the different brain regions of instrumental mice (Extended Data Fig. 1E). Among the FFAs that increased following instrumental conditioning, saturated FFAs, particularly stearic acid (C18:0), palmitic acid (C16:0), myristic acid (C14:0), and hexanoic acid (C6:0) constituted the majority, while polyunsaturated FFAs such as arachidonic acid (C20:4), and monounsaturated FFAs such as oleic acid (C18:1), palmitoleic acid (C16:1), and nervonic acid (C24:1) represented the predominant unsaturated FFAs across all brain regions (Extended Data Fig. 1E). We also observed a significant reduction in octanoic acid (C8:0), llignoceric acid (C24:0), and erucic acid (C22:1) across the different brain regions (Extended Data Fig. 1E). In agreement with previous results obtained in auditory fear conditioned rats (Wallis *et al*., 2021), our data revealed that instrumental conditioning drives an overall increase in FFAs levels across the key brain regions, with the most significant changes seen in saturated FFAs.

### *DDHD2^-/-^* drives progressive decline in spatial memory, motor function, and explorative behaviors

To explore the impact of the DDHD2 knockout, we carried out a longitudinal assessment of the age of onset and progression of various neuromuscular memory deficits, in *DDHD2^-/-^* mice (genetically engineered to knock out the *DDHD2* gene) in comparison to their wild-type *DDHD2^+/+^* litter mates, monthly from 3mo to 12mo as detailed in **Methods**. Motor coordination and strength were assessed using rotarod and grip strength tests respectively. The onset of impairment in motor coordination was observed at 5 months in *DDHD2^-/-^* animals and from this time point progressively declined throughout the period of the study (Extended Data Fig. 2A). In comparison to the wild-type animals, *DDHD2^-/-^* mice demonstrated a progressive decline in motor strength, with an onset at 6 months (Extended Data Fig. 2B). These results suggest that the *DDHD2* KO drives a progressive decline in motor function in mice beginning from 5mo, which further corroborates previous reports suggesting that the ablation of the *DDHD2* gene elicits the manifestation of HSP and its associated neuromotor dysfunction symptoms (Blackstone, 2018; Murala *et al*., 2021).

We adapted a longitudinal novel object location (NOL) test to investigate short-term spatial memory impairment. The NOL behavioural paradigm relies on the innate instincts of mice to explore novel locations more than familiar ones (Blackmore *et al*, 2022). Results demonstrated a significant decrease in the time that *DDHD2^-/-^* mice spent exploring the NOL compared to the wild-type mice from 3mo onwards, until the termination of the study (Extended Data Fig. 2C). This suggests that disruption of the *DDHD2* gene drives age-dependent impacts on spatial memory.

Using automated activity monitoring, we also longitudinally assessed open field locomotor performance, alongside explorative and vigilance activity. 3mo *DDHD2^-/-^* mice exhibited a significant decrease in vertical time, vertical counts, jump time, and jump counts when compared to *DDHD2^+/+^* mice. The observed decrease was progressive and more significant in 12mo mice (Extended Data Fig. 2D-H). The progressive decrease in these observations suggests impaired vigilance, exploration, and escape response. There was no significant difference in the resting time and ambulatory distance of *DDHD2^+/+^ and DDHD2^-/-^* mice. Taken together the results suggest that *DDHD2^-/-^*drives progressive age-dependent alteration in mice behaviour and consequently DDHD2 is critically involved in spatial memory via production of saturated FFAs.

### Disruption of the *DDHD2* gene drives age-dependent reduction in long-term memory performance and saturated FFAs levels (including myristic acid)

The mammalian PLA1 family of enzymes consists of both intracellular and extracellular isoforms, all of which contain a DDHD domain. Among the 13 mammalian PLA1 isoforms, only three are intracellular: iPLA1α (also known as DDHD1 and phosphatidic acid-preferring PLA1; PA-PLA1), iPLA1β (also known as p125), and iPLA1γ (also known as DDHD2) (Inoue *et al*., 2012). Activation of the DDHD2 domain has previously been linked with an increased generation of reactive oxygen species and apoptosis (Maruyama *et al*, 2018), mitochondrial dysfunction and mitochondrial phospholipid remodelling (Yadav & Rajasekharan, 2016), lipid trafficking, and signalling (Baba *et al*, 2014; Lev, 2004). DDHD2 is thought to utilise phosphatidic acid (PA) as its preferred substrate, and by doing so generates lysophosphatidic acids (LPAs) and saturated FFAs (Baba *et al*., 2014; Higgs & Glomset, 1996). We therefore sought to elucidate the effect of knocking out the *DDHD2* gene, which codes for an endogenous PLA1 that has a role in the generation of FFAs during longer-lasting memory acquisition *in vivo* (Inloes *et al*., 2014; Inloes *et al*, 2018). Instrumental conditioning was carried out using separate cohorts of *DDHD2^-/-^* vs *DDHD2^+/+^* litter mates at 3mo and 12mo. Data obtained from instrumentally conditioned *DDHD2^-/-^* mice exhibited a trend towards memory impairment at 3mo (asymptomatic young animals; Extended Data Fig. 1A-C), and a statistically significant memory deficit at 12mo (symptomatic animals; Fig. 1B)

Using FFAST LC-MS/MS (Narayana *et al*., 2015) as described above, we quantified the abundance of 19 targeted FFA species from 5 brain regions of instrumentally conditioned *DDHD2^+/+^* and *DDHD2^-/-^* mice at 3mo and at 12mo, and by doing so uncovered a significant decrease in the levels of most FFAs across all targeted brain regions in *DDHD2^-/-^* mice (Extended Data Fig 3A-E). Principal component analysis (PCA) was used to dimensionally reduce the average FFA profiles from 19 dimensions to 3, revealed that the FFA profiles from instrumentally conditioned *DDHD2^+/+^*and *DDHD2^-/-^* mouse cohorts cluster separately from their respective control mouse cohorts, at both 3mo and at 12mo (Extended Data Fig. 4A-C). Examination of the underlying components using scatter plots of absolute change vs fold change showed that the observed clustering was predominantly driven by saturated FFAs such as myristic acid (C14:0), palmitic acid (C16:0), and stearic acid (C18:0), and to a lesser extent, mono-unsaturated FFAs docosahexaenoic acid (C22:6), linolenic acid (C18:3), and arachidonic acid (C20:4), which decreased the most in instrumentally conditioned *DDHD2^-/-^*mice (Fig. 1C-F). Overall, our data suggests that DDHD2 substantially contributes to lipid metabolism through the generation of saturated FFAs, and that its ablation triggers both a major reduction in the concentration of these FFAs across different regions of the brain, resulting in a blunted response following memory acquisition tasks.

### STXBP1 is a major DDHD2 binding protein

The mechanism by which DDHD2 activity mediates memory consolidation in fear conditioning (Wallis *et al*., 2021), instrumental and spatial memory testing is currently unknown. The fact that saturated FFA increases are activity-dependent makes it likely that the activity of this enzyme is localized at the synapse. Although it is known that DDHD2 is expressed in neurons (Inloes *et al*., 2014), the mechanisms through which it is trafficked to the synaptic compartment where it modifies the lipid landscape are unclear. We hypothesized that the production of FFAs is likely associated with synaptic activity, and regulated by a resident synaptic protein. To investigate this, we carried out a series of co-immunoprecipitation pull-down assays from neurosecretory Pheochromocytoma (PC12) cells to identify molecules that may potentially interact with DDHD2. Untargeted high-resolution tandem mass spectrometry (HRMS) analysis of peptide digests from a DDHD2 pull down identified 150 proteins at a 1% false discovery rate (Extended Data Fig. 5, Table 4 and 5 for a full list of identified proteins). The number of proteotypic peptides with an identification confidence score above the 95% acceptance threshold, and the level of coverage these peptides have with the sequence of the identified protein were used as an approximate measure of the abundance of each protein that was identified. Reassuringly, using this metric, DDHD2 was the most abundant protein identified in the protein extracts – with 206 unique proteotypic peptides identified, covering 80.4% of the DDHD2 sequence (Uniprot accession: D3ZJ91). The second most abundant protein identified was Syntaxin-binding protein 1 (STXBP1, also known as Munc18-1; Uniprot accession: P61765) with 94 peptides identified, covering 76.4% of the protein sequence. This indicates that STXBP1 is a primary binding partner of DDHD2. STXBP1 is a synaptic protein that is critically involved in the priming of secretory and synaptic vesicles (Deak *et al*, 2009; Kasula *et al*) and is an essential component of the presynaptic neurotransmitter release machinery –with its knockout leading to a complete loss of neurotransmitter release and perinatal death (Martin *et al*, 2013; Verhage *et al*, 2000).

To explore this further, we performed a functional association analysis by searching the list of identified proteins against the Search Tool for Retrieval of Interacting Genes/Proteins (STRING) database (Szklarczyk *et al*, 2021).This revealed that among the proteins that were co-purified with DDHD2 under a resting state (unstimulated), the primary interactome of STXBP1 consisted of six significant interactions including Rab3c, Rab10, Rab5c, Nefm, Ppp1r9b and Sec22b (Fig. 2A). The primary role of these proteins involves vesicle and membrane organisation, suggesting that STXBP1 may be acting as an interface between DDHD2 and membrane organising structures. Gene ontology cellular component analysis revealed other proteins co-purifying with DDHD2 to a lesser degree (Extended Data Fig. 5). To further validate the interaction between STXBP1 and DDHD2, a targeted multiple reaction monitoring mass spectrometry (MRM-MS) assay specific for sevenproteotypic peptides for STXBP1, and for four proteotypic peptides for DDHD2, was employed. This assay was applied to the DDHD2 pull down, providing a parallel mass spectrometric measurement confirming the HRMS identifications of both DDHD2 and STXBP1, and was also applied to a STXBP1 pull-down from PC12 cells, which confirmed the cognate presence of DDHD2 (Extended Data fig. 5). This suggests that DDHD2 and STXBP1 undergo a biologically significant interaction. To assess whether the binding between DDHD2 and STXBP1 was altered in response to cellular stimulation, the targeted MRM-MS assay was employed to quantitatively assess the relative amount of STXBP1 that co-purified with DDHD2 in response to high K^+^ stimulation prior to pull down This resulted in a 2.3-fold significant increase in STXBP1 associated with DDHD2 immediately after high potassium depolarisation compared to unstimulated controls, indicating that an activity dependent increase in binding was occurring (Fig. 2B). To further confirm this binding, we used gene-edited neurosecretory cells lacking STXBP1/2 (PC12-DKO (Kasula *et al*.)), which we transfected with STXBP1-emGFP. Cells were stimulated using high K^+^ and solubilized for pull down via GFP-trap. We further confirmed that DDHD2 was detected in the pull-down fraction and that it was largely increased in response to stimulation (Fig. 2C). Conversely, we expressed GFP-DDHD2 and performed GFP-Trap, and detected STXBP1 by western blotting, which was significantly increased following stimulation (Fig. 2D).

**Fig. 2.**
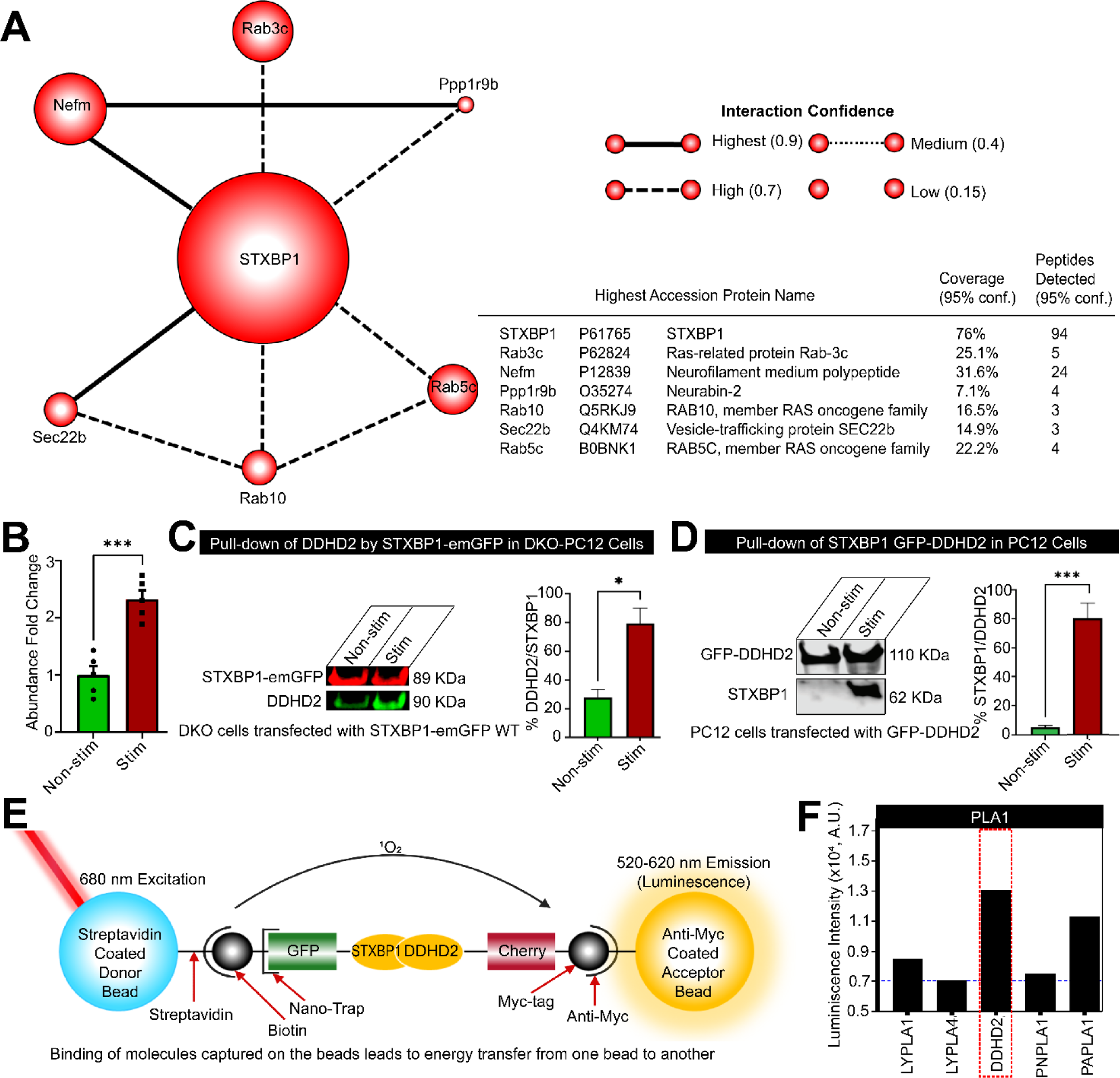
Identification of STXBP1 (Munc18-1) as a major DDHD2 interactor. **A** STXBP1 and its direct interaction partners identified using STRING (https://string-db.org/) functional association analysis of proteins co-precipitating with DDHD2 as determined by untargeted high resolution tandem mass spectrometry (HRMS). The relative size of each protein icon represents the percentage of sequence coverage with high quality (>95% identification confidence score) proteotypic peptides detected by HRMS. The thickness and hatching of the lines connecting each protein represents the confidence score of the strength of the interaction as determined by the STRING knowledge base matching algorithm. **B** Targeted MRM mass spectrometry showing relative abundance of STXBP1 peptides co-precipitated with DDHD2 in high vs low K+ stimulated cultures as described in **A**. **C** Binding of STXBP1 with DDHD2. Immunoblot of STXBP1-emGFP (89 KDa, red) pulldown of DDHD2 (90 KDa, green) and **D** GFP-DDHD2 (110 KDa) pulldown of STXBP1 (62 KDa) in non-stimulated and BaCl_2_-stimulated PC12 cells. Densitometric quantifications of the immunoblots are shown as percentage of DDHD2 to STXBP1 and STXBP1 to DDHD2 respectively. Results are shown as mean ± SEM from four independent experiments. The significance of the change between the different experimental conditions as determined by two-tailed unpaired Student’s t test, * p<0.05, *** p<0.001. The significance of the change between the different experimental conditions as determined by two-tailed unpaired student’s t test. **E** Schematic of proximity based ALPHAScreen analysis using tagged *in vitro* translated proteins to illustrate STXBP1^WT^ interactions with phospholipases. The biotin-coupled GFP-nanotrap recruits the GFP tagged proteins (STXBP1^WT^) and binds with streptavidin-coated donor beads. Myc-tagged proteins (phospholipases, all expressed at the same concentration) bind to anti-myc coated acceptor beads. The donor bead bound to STXBP1^WT^ releases singlet oxygen (^1^O_2_) upon illumination at 680 nm, which only diffuses 200 nm in solution. Upon reaching an acceptor, ^1^O_2_ reacts with derivatives in the acceptor bead containing phospholipases and luminescence is detected at 520-620 nm. The luminescence observed indicate the relative strength of the binding. **F** ALPHAScreen STXBP1^WT^ interactions with phospholipases. The strong interaction of STXBP1 and DDHD2 is highlighted with a red box.

Finally, we investigated whether DDHD2 directly binds to STXBP1 *in vitro,* using tagged proteins in the proximity-based Amplified Luminescent Proximity Homogeneous Assay Screen (ALPHAScreen) (Martin *et al*., 2013; Sierecki *et al*, 2013). In this assay, streptavidin-coated donor beads bind a biotin-coupled GFP-nanotrap that recruits GFP-tagged STXBP1^WT^ protein. The acceptor beads coated with anti-myc antibody, bind to myc-tagged DDHD2 and other myc-tagged PLA1 isoforms (Fig. 2E). Protein-protein interactions are detected via energy transfer luminescence. The ALPHAScreen assay confirmed that STXBP1 directly interacts with DDHD2, and to a lesser extent, with PAPLA1, PNPLA1, and LYPLA1 2 isoforms (Fig. 2E-F).

### STXBP1 controls DDHD2 transport to the plasma membrane

Having demonstrated that STXBP1 binds to DDHD2, we next sought to investigate the functional significance of this interaction. STXBP1 was originally described and named due to its significant interaction with syntaxin (STX), with several studies showing that STXBP1 binds to STX1A to mediate its transport to the plasma membrane of neurosecretory cells (Martin *et al*., 2013; Rickman *et al*, 2007), and to mediate its organisation and engagement into the SNARE complex during neuroexocytosis. We therefore investigated whether STXBP1 is responsible for the transport of DDHD2 to the plasma membrane via a similar mechanism.

We performed immunostaining of neurosecretory PC12 cells and found that DDHD2 was largely localized to the periphery of the cells, suggestive of plasma membrane localisation (Fig. 3A-D). Further, DDHD2 plasma membrane localisation was similar to that of STX1A (Fig. 3E-H). We used gene-edited PC12 cells genetically-engineered to knock out STXBP1/2 (Kasula *et al*.) to investigate the role of STXBP1 in the transport of DDHD2 to the plasma membrane and observed that in the absence of STXBP1/2, both DDHD2 and STX1A were mislocalised away from the plasma membrane (Fig. 3I-K). Analysis of DDHD2 with STX1A in PC12 cells showed high level of co-localization on the plasma membrane which was much reduced and mislocalised in DKO PC12 cells (Fig. 3L). Re-introducing GFP-tagged wild-type STXBP1 into DKO PC12 cells rescued the plasma membrane localization of DDHD2 and STX1A (Fig. 3M-P), indicating that STXBP1 has a central role in the transport of DDHD2 to the plasma membrane. We checked whether STXBP1 could also control the expression levels of DDHD2 *in vitro*, however, the relative protein expression levels were not significantly different between PC12 cells and STXBP1/2 DKO PC12 cells when assessed by western blot assay and PCR (Extended Data Fig. 6A-B). Remarkably, hippocampal neurons from DDHD2^-/-^ animals exhibited instances of arrested secretory vesicles suggestive of a transport defect in the ER-Golgi intermediate compartment (ERGIC; Extended Data Fig. 7). Together, these results suggest a novel role for STXBP1 in transporting DDHD2 to the plasma membrane, and in the subsequent release of saturated FFAs during stimulated exocytosis.

**Fig 3.**
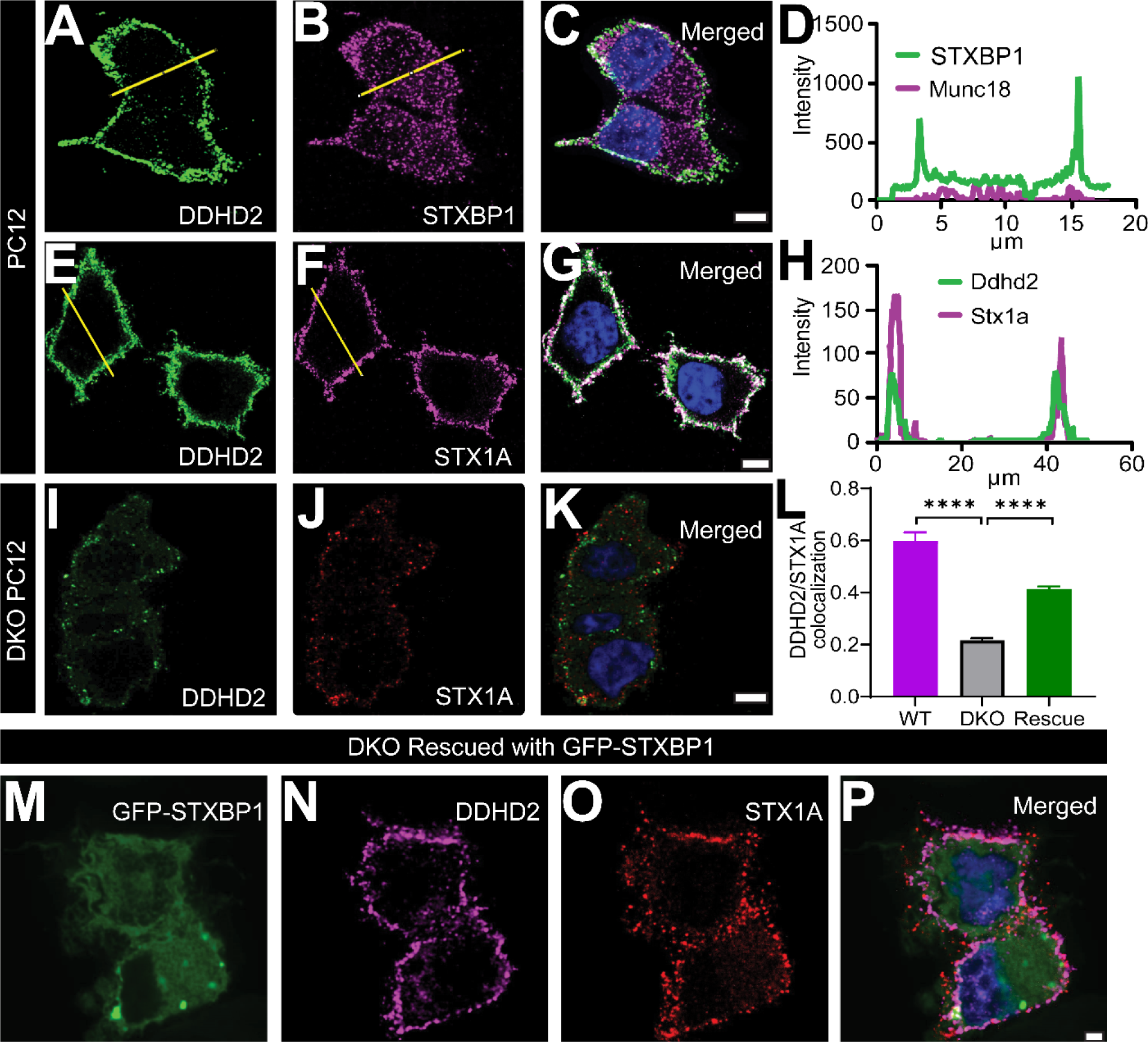
STXBP1 (Munc18-1) plays a key role in the transport of DDHD2 to the plasma membrane. Representative image of BaCl_2_-stimulated PC12 cells immunostained for **A** endogenous DDHD2 and **B** endogenous STXBP1. The merged image of the channels is shown in **C**, and the fluorescence intensity profile of DDHD2 (green) and STXBP1 (magenta) from the indicated yellow line is shown in **D**. Representative images of BaCl_2_-stimulated PC12 cells immunostained for **e** endogenous DDHD2 and **F** endogenous STX1A. The merged image of the channels is shown in **G**, and the fluorescence intensity profile of DDHD2 (green) and STX1A (magenta) from the indicated yellow line is shown in **H**. Representative images of BaCl_2_-stimulated STXBP1/2 double knockout (DKO) PC12 cells stained for **I** endogenous DDHD2 and **J** endogenous STX1A. The merged image of the channels is shown in **K**, and the quantification of the DDHD2 co-localization with STX1A in PC12 cells, DKO PC12 cells, and DKO PC12 cells following rescue with GFP-STXBP1 **M-P** is shown in **L**. Scale bars are 5 µm. **** p<0.0001.

### STXBP1 binding to DDHD2 controls FFA levels

To assess the potential of STXBP1 in controlling the activity-dependent increase in FFAs from neurosecretory cells, we used FFAST to compare the FFA response to secretagogue stimulation (high K^+^) in wild type (WT) PC12 cells and gene-edited neurosecretory cells lacking STXBP1/2 (DKO PC12). Our results showed that stimulation of PC12 cells led to a significant increase in predominantly saturated FFAs, particularly C14:0, C16:0, and C18:0 (Fig. 4A) as previously detected in chromaffin cells (Narayana *et al*., 2015). Conversely, in DKO PC12 cells lacking STXBP1 and 2, the basal (unstimulated) FFA levels were significantly reduced compared to WT PC12 cells, suggesting that STXBP1 is required for maintaining basal FFA levels in PC12 cells (Fig. 4B). Additionally, the activity-dependent increase in saturated FFAs was abolished in stimulated DKO PC12 cells, indicating that STXBP1 controls the production of saturated FFAs in response to secretagogue stimulation (Fig. 5A). As previously described for Munc18-1/2 double-knockdown neurosecretory cells (Han *et al*, 2010; Han *et al*, 2009), DKO PC12 cells were also unable to secrete exogenous neuropeptide-Y (NPY) in response to depolarising stimulus in a NPY-human placental alkaline phosphatase (NPY-hPLAP) release assay (Extended Data Fig. 6C). Taken together, both baseline lipid homeostasis and activity-dependent FFA generation correlated with functional exocytosis release, and are likely controlled by STXBP1 (Fig. 4A, Extended Data Fig. 6C).

**Fig 4.**
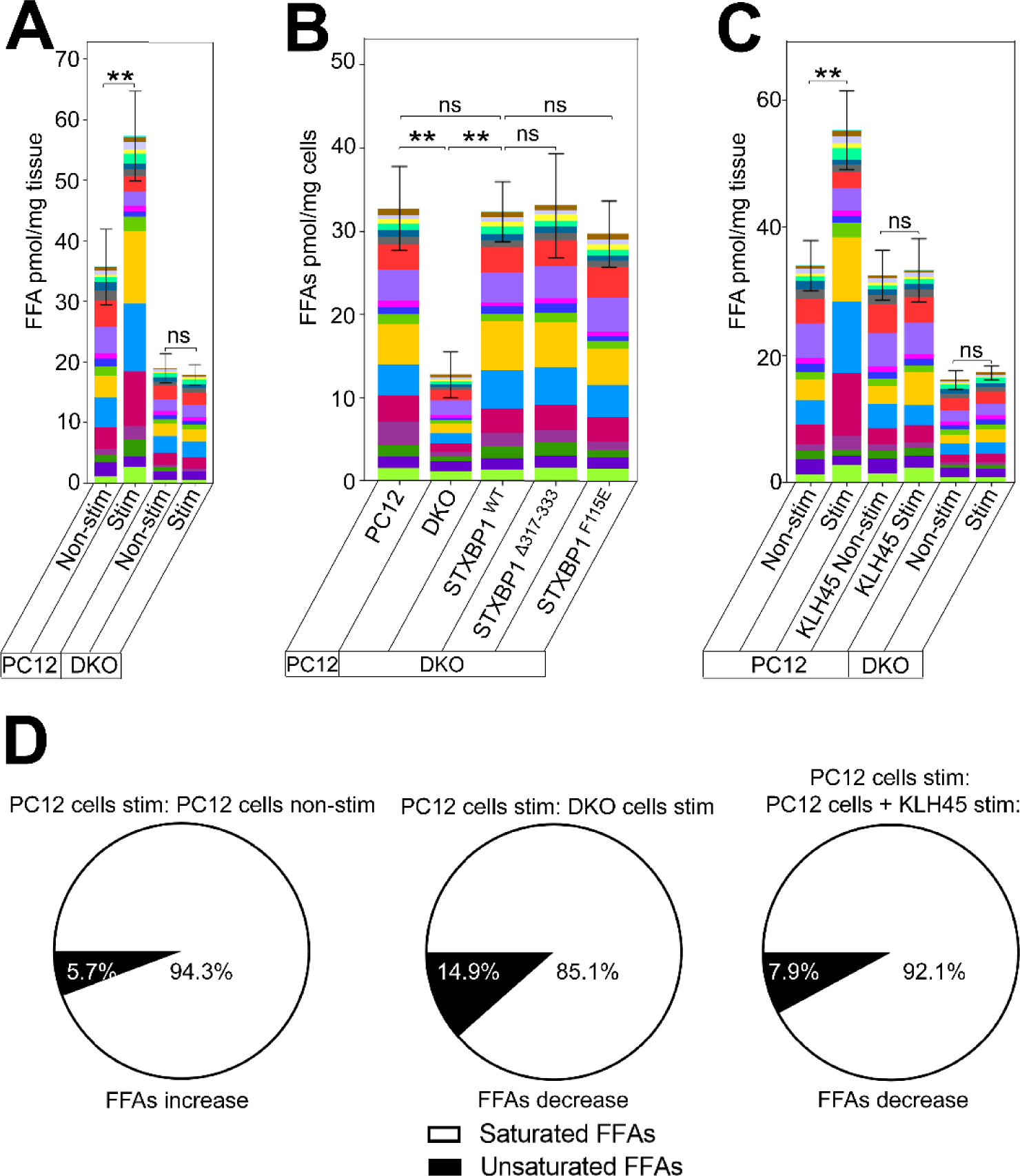
STXBP1 (Munc18-1) controls the generation of saturated FFAs *in vitro* and *in vivo*. FFA abundance determined using FFAST, as detailed in Fig. 1**. A** Stacked bar plot showing profiles of FFAs in PC12 cells and STXBP1/2 double knockout (DKO) PC12 cells following stimulation. PC12 and DKO cells were stimulated (Stim) by depolarization for 15 min in high K^+^ (60 mM) buffer. Control unstimulated (Non-stim) cells were treated for 15 min in low K^+^ (2 mM) buffer. The significance of the change in average FFA abundance between PC12 Non-stim/Stime and DKO Non-stim/Stim as determined by t-test with Holm-Sidak *post-hoc* correction is indicated by asterisks ** p<0.01. ns = not significant. **B** Basal FFAs levels in PC12 cells, STXBP1/2 DKO PC12 cells, DKO cells transfected with STXBP1^WT^, DKO cells transfected with STXBP1^Δ317-333^ (Loop mutant), and DKO cells transfected with STXBP1^F115E^ (hydrophobic pocket mutant) observed across 3 replicates (pmol/mg tissue). **C** Profile of FFA responses to stimulation in PC12 cells following pharmacological inhibition of DDHD2 using KLH45 (-/+ stimulation), and in DKO PC12 cells. **D** Pie charts showing the percentage change of saturated vs unsaturated FFAs in different conditions. The significance of the change in FFA abundance between the different experimental conditions as determined by one-way ANOVA with Holm-Sidak *post-hoc* correction is indicated by asterisks ** p<0.01, ns = not significant. **A, B, C** error bars represent the cumulative standard error of the mean (SEM) for all groups and parameters.

**Fig. 5.**
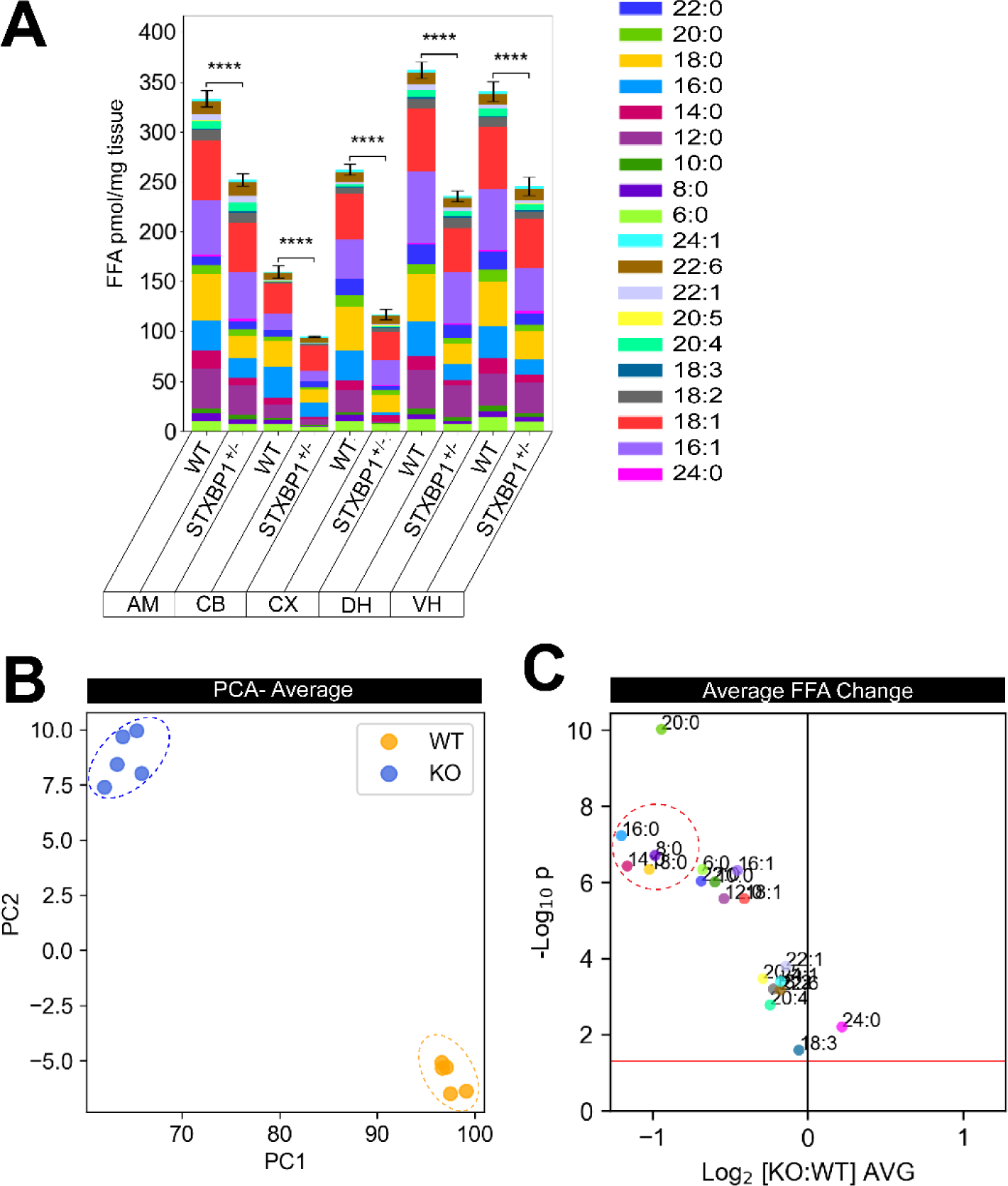
FFA response in STXBP1 haploinsufficiency mice brain. **A** Bar plots showing the total FFA levels across different brain regions (CB; cerebellum, CX; cortex, DH; dorsal hippocampus, VH; ventral hippocampus, AM; amygdala) of wild type (WT) versus STXBP1^+/-^ (KO) mice (n=5). The significance of the change in FFA abundance between the different experimental conditions as determined by one-way ANOVA with Holm-Sidak *post-hoc* correction is indicated by asterisks, **** p<0.0001. **B** Principal component analysis of the average FFA profile across the brain. **C** Volcano plot showing average response across the brain of individual FFAs in WT versus KO mice. Each dot on the volcano plot represents the average change in abundance of a single analyte across 5 measured brain regions. Analytes below the red line represent those whose change in abundance was not statistically significant (two-tailed *t*-test *p* > 0.05). Error bars represent the cumulative standard error of the mean (SEM) for all groups and parameters.

### STXBP1 controls the generation of saturated FFAs

We next sought to establish whether the FFA lipid profile can be rescued upon re-expression of WT STXBP1 in STXBP1/2 DKO PC12 cells. Our results revealed that re-expression of WT STXBP1 WT in these DKO PC12 cells completely rescued the FFA basal levels, demonstrating that STXBP1 is required to control DDHD2-induced remodelling of the neurosecretory membrane (Fig. 4B). Since STXBP1 is a key regulator of vesicular priming in both secretory and synaptic vesicles (Deak *et al*., 2009; Kasula *et al*, 2016), we wanted to assess whether STXBP1 missense mutants with altered priming and STX1A binding were similarly able to rescue the FFA profile. Both the re-expression of STXBP1^Δ317-333^, a priming deficient loop mutant that prevents the opening of STX1A and SNARE assembly (Kasula *et al*., 2016; Martin *et al*., 2013), and STXBP1^F115E^, a hydrophobic pocket mutant (HPM) that ablates the binding of the SNARE complex with STXBP1 (Han *et al*., 2010; Han *et al*., 2011; Malintan *et al*, 2009), was able to restore the levels of saturated FFAs (Fig. 4B). This demonstrated that DDHD2 transport to the plasma membrane is underpinned by STXBP1 and is independent of STXBP1’s function in vesicular priming.

To confirm that DDHD2 is also required during the activity dependent FFA response, we pre-treated PC12 cells with a pharmacological inhibitor of DDHD2, KLH45, prior to stimulated exocytosis and FFAST analysis. As we hypothesised, acute pharmacological inhibition of DDHD2 abolished the activity-dependent increase in saturated FFAs (Fig. 4C-D). Comparison of the FFA response in PC12 cells, STXBP1/2 DKO PC12 cells, and PC12 cells treated with KLH45 showed that the activity-dependent FFA changes that were observed across the different conditions were largely driven by saturated FFAs (Fig. 4D).

### STXBP1 haploinsufficiency leads to FFA deficits in the brain

STXBP1 is an essential component of the presynaptic neurotransmitter release machinery and its knockout leads to a complete loss of neurotransmitter release and perinatal death (Verhage *et al*., 2000). Importantly, *de novo* pathogenic mutations of this gene (either splice site variants, nonsense, frameshifts, or deletions) lead to neonatal mortality associated with pronounced neurodegeneration (Heeroma *et al*, 2004; Verhage *et al*., 2000). Further, alteration in the expression of STXBP1 caused by a range of human variants in the *STXBP1* gene leads to developmental epileptic encephalopathies (DEE) – a set of neurodevelopmental conditions associated with intellectual disability (STXBP1-DEE) (Abramov *et al*, 2021; Chen *et al*., 2020; Guiberson *et al*, 2018; Miyamoto *et al*, 2017). STXBP1 encephalopathy encompasses a range of neurodevelopmental conditions including autism, intellectual disability (mental retardation), cognitive impairment, and movement disorder (Chai *et al*, 2016; Lanoue *et al*, 2019; Saitsu *et al*, 2010; Stamberger *et al*, 2016; Tavyev Asher & Scaglia, 2012), and is thought to stem from haploinsufficiency altering the levels of functional STXBP1 protein and gain of toxic function (Lanoue *et al*., 2019). Having shown herein that STXBP1 controls the production of saturated FFAs critical for memory acquisition (Wallis *et al*., 2021), we hypothesized that haploinsufficiency would lead to reduced saturated FFA production. We used a genetically distinct haploinsufficient mouse model (*STXBP1*^+/-^) showing a 40–50% decrease in the levels of STXBP1 protein, which recapitulates the cognitive, motor, seizure, and psychiatric phenotypic hallmarks of human encephalopathy (Abramov *et al*., 2021). As anticipated, FFAST LC-MS/MS analysis of 19 FFAs from 5 different brain regions from the *STXBP1*^+/-^ mouse revealed a significant decrease in the total FFA levels in heterozygous mice compared to WT animals that have two copies of STXBP1 (Fig. 5A, Extended Data Fig. 8A-B). Further analysis of this data using principle component analysis (PCA) and partial least square discriminant analysis (PLSDA) of the average FFA profile across the brain showed that the heterozygous animals have a distinct FFA profile (Fig. 5B, Extended Data Fig. 8C), largely driven by saturated FFAs (C12:0, C14:0, C16:0 and C18:0) and to a lesser extent monounsaturated FFAs (C16:1 and C18:1) which were significantly decreased compared to WT animals (Fig. 5A, C, Extended Data Fig. 8A, B). Together, our results point to a novel role for STXBP1-DDHD2 interaction in the regulation of the saturated FFA landscape in the brain.

#### Discussion

This study utilised a multidisciplinary approach to uncover the mechanisms underlying changes in the FFA landscape of the brain that takes place during memory formation. We used targeted FFAST lipidomics coupled with a reward-based long-term memory paradigm, to demonstrate that the DDHD2 isoform of PLA1 is critical for driving saturated FFA responses to memory acquisition, with its ablation leading to the progressive decline in memory as well as neuromuscular performance. We also report a novel interaction between DDHD2 and STXBP1 – a key regulator of the exocytic machinery. Further, we demonstrate that STXBP1 controls the transport of DDHD2 to the plasma membrane and the subsequent generation of saturated FFAs (particularly myristic acid) during stimulated neuroexocytosis. Together this data demonstrates that DDHD2 regulates FFA changes that are associated with memory via interaction with STXBP1.

Consistent with previous studies which showed that saturated FFAs predominate the FFA response in both *in vitro* stimulation of neuronal cultures (Narayana *et al*., 2015) and during fear memory acquisition *in vivo* (Wallis *et al*., 2021), our observation that instrumental conditioning also induced changes in the saturated FFA landscape of the brain indicates that regardless of the paradigm used, localised activity-dependent increases in saturated FFAs in the brain is a general feature of memory. Due to the long-term nature of instrumental conditioning, a greater intensity in FFA responses across all the assessed brain regions was observed, and in contrast to fear conditioning where the highest FFA response was elicited in the amygdala, we uncovered the highest FFA response in the dorsal hippocampus and cortex, signifying a brain region-specific FFA response to reward-based memory. These findings can be attributed to the involvement of the dorsal hippocampus, prefrontal cortex, and the posterior dorsomedial striatum which form a critical hub for the complicated set of circuits that network together to encode reward-based memory in the brain (Balleine, 2019; Balleine & O’Doherty, 2010), and it is expected that FFA changes indicative of synaptic plasticity would be concentrated in these areas. Although the contribution of the cerebellum to non-motor function has been suggested (Timmann & Daum, 2007), the lower FFA response observed in this brain region may be ascribed to its primary involvement in motor processes (Statton *et al*, 2018) and consequently, its lower relevance to synaptic plasticity in response to instrumental conditioning. Although poly unsaturated fatty acids (PUFAs) such as AA, DHA, and eicosapentaenoic acid (EPA), previously associated with membrane fluidity (Fukaya *et al*, 2007), neuronal signalling and SNARE complex formation (Falomir-Lockhart *et al*., 2019; Garcia-Martinez *et al*, 2018; Rickman & Davletov, 2005), as well as memory (Inoue *et al*, 2019), also changed in our study, the overall response was dominated by saturated FFAs. The significant reduction in octanoic acid (C8:0), lignoceric acid (C24:0), and erucic acid (C22:1) which we observed across the different brain regions may be attributed to their energy-contingent usage or their metabolism as a precursor for generating other FFAs in response to memory acquisition (Andersen *et al*, 2021).

Our findings further suggest that the DDHD2 isoform of PLA1 may be activated during these processes, and supports the idea that PLA1 plays a critical role in memory formation via the regulation of saturated FFA generation and dynamically modifying the phospholipid landscape of the brain (Higgs & Glomset, 1996; Inloes *et al*., 2014). The substrate-product relationship of DDHD2 is interesting as relatively few phospholipids exhibit a C14:0 on the *sn-1* position. Most of the fatty acyl chains are either C16:0 or C18:0 and although they also increase in response to conditioning, their levels are consistently lower compared to the amount of C14:0 generated. It is therefore reasonable to speculate that DDHD2 exhibits some level of selectivity for the C14:0 acyl chain phospholipids. Additional work will be needed to address this issue. Further, a decrease in unsaturated FFAs was detected in *DDHD2* knockout mice which could be attributed to the presence of non-canonical phospholipid substrates that do not have a saturated *sn-1* and an unsaturated *sn-2* acyl chain configuration (Thomas *et al*, 2006; Wang & Hsu, 2022). Alternatively, it may also stem from the reduced specificity of the PLA1 enzyme in cleaving both *sn-1* and *sn-2* positions of the fatty acyl chain of the phospholipid substrates. In line with this hypothesis, genetic dysregulation affecting DDHD2 expression (Inloes *et al*., 2014) and its localisation, greatly impacts saturated FFA levels and memory acquisition. Our findings therefore add weight to the theory that saturated FFAs are critical in memory formation and contribute to the mechanism(s) underpinning the retention of memory and cognitive function (Zamzow *et al*, 2019).

Beyond establishing a link between saturated FFA levels and brain function, our study provides relevant insights into the onset and progressive impairment of memory performance in *DDHD2^-/-^*mice. The discovery of a trend in memory impairment coupled with changes in the brain FFA landscape of otherwise asymptomatic 3mo mice, prior to the onset of a decline in motor function at 5mo, suggests that impairment in memory function precedes the onset of motor dysfunction in *DDHD2^-/-^* mice. The impairment of memory performance associated with the DDHD2 knockout reported in our study could, at least in part, explain the intellectual disability associated with HSP, and recent studies showing that ablation of the *DDHD2* gene alters neural processing (Inloes *et al*., 2014; Joensuu *et al*., 2020; Richmond & Smith, 2011) also strongly advocate for the significant impact of saturated FFAs in synaptic function and plasticity.

Our observed decline in motor function corroborates earlier reports associating disruption of the *DDHD2* gene with classical signs of slowly progressing spastic paraparesis characterised by weakness, hyperreflexia, neuromotor dysfunction predominantly affecting the lower limbs, and ultimately aberrant gait (Blackstone, 2018; Inloes *et al*., 2014; Inloes *et al*., 2018; Joensuu *et al*., 2020; Parodi *et al*, 2017). The observation that deficits in motor coordination preceded decline in motor strength suggests that the premotor cortex, which is responsible for some aspects of motor control including preparation for movement and sensory guidance of movement, may be impacted more significantly or earlier than the primary motor cortex, which mainly contributes to the generation of neural impulses that control the execution of movement (Cisek & Kalaska, 2005). Hence, motor coordination may be a preferable diagnostic test for the early detection of HSP. The observed motor deficit may be attributable to changes in the FFA landscape of the cerebellum in genetically ablated *DDHD2* animals (Janssen *et al*, 2015).

Another important aspect of our study was the discovery of a novel function for STXBP1 in transporting DDHD2 to the plasma membrane for the generation of saturated FFAs which occurs independently of pre-exocytic vesicular priming. This is in line with the previously reported transport role of Munc18 as a chaperone for proteins such as STX. We describe a mechanism by which STXBP1 can allow targeting of DDHD2 to the presynapse, where it dynamically modifies the lipid landscape to generate saturated FFA metabolites during memory formation. Consistent with a major role of saturated FFAs in learning and memory, we also found that haploinsufficient STXBP1 mice have much reduced saturated FFAs which may potentially explain their poor cognitive performances (Chen *et al*., 2020) and that of STXBP1 encephalopathy patients (Abramov *et al*., 2021; Lanoue *et al*., 2019; Stamberger *et al*., 2016). This finding also suggests the possible involvement of saturated FFAs in the pathophysiological mechanisms underlying DEE resulting from STXBP1 mutations. Further, previous studies have shown that mutations in the *DDHD2* gene regulate Golgi-/ER membrane trafficking and further support its critical role in essential cellular processes associated with memory function. The observation of arrested vesicles along the Golgi-/ER interface of hippocampal neurons from *DDHD2^-/-^* mice is suggestive of a transport defect which may also be contributing to the memory deficit observed in these animals. This result corroborates previous reports which used electron microscopy to detect an accumulation of lipid droplets and multilamellar bodies which can alter autophagy-lysosome function, and render the cell more susceptible to apoptosis, subsequently resulting in memory dysfunction (Garcia-Sanz *et al*, 2018).

Although FFAs can mediate the process of memory consolidation via several mechanisms including the modulation of membrane properties, post-translational targeting of proteins to interact with membranes and other proteins, or via other lipid signalling pathways, how these saturated FFAs affect synaptic function is currently unknown. We hypothesize that synaptic protein acylation occurring via acyl-CoA intermediates, is a key player in this process (Seo *et al*, 2022). Considering that myristic and palmitic acids are highly increased in response to memory acquisition, protein lipidation could be playing a role in the establishment of synaptic plasticity. More work is needed to assess this important question.

Together, our findings demonstrate for the first time that the interaction between DDHD2 and STXBP1 is critical for long-term memory by regulating the activity-dependent generation of FFAs at the synapse, and dynamically modifying the lipid landscape. Consequently, DDHD2 may be an important pharmacological target for regulating key FFA signalling pathways, and a better understanding of the DDHD2-regulated lipid pathways may offer novel insights into the mechanism and therapeutic strategies for the cognitive deficits associated with ageing and neurodegenerative disease.

## Material and Methods

### Ethical considerations and animals

For all experimental procedures, the care and use of animals was carried out in line with the protocols approved by the Animal Ethics Committee of The University of Queensland (2017/AE000497, 2018/AE000508, 2021/AE000971, 2020/AE000352 and 2022/AE00073) and by the Institutional Animal Care and Use Committee at Baylor College of Medicine (protocol AN-6544).

### Key resources

HPLC / analytical grade reagents were used throughout. 1,1-carbonydiimidiazole, triethylamine, iodomethane, iodomethane-d3, iodoethane, iodoethane-d5, iodopropane, formic acid, citric acid, methanol, disodium hydrogen phosphate, chloroform, ammonium formate, acetonitrile, 1-butanol, and analytical standards for saturated and unsaturated fatty acids were purchased from Sigma-Aldrich. All lipid extractions were performed in 2 mL polypropylene LoBind safe-lock tubes (Eppendorf).

Cell culture reagents were purchased from Life Technologies. Mouse anti-STXBP1 antibody was purchased from BD Biosciences. pCMV-STXBP1-emGFP, pCMV-STXBP1, NPY-hPLAP, and NPY-mCherry were prepared as previously described (Arunachalam *et al*., 2008; Martin *et al*., 2013; Tomatis *et al*, 2013). STXBP1^Δ317-333^ was made using the quick-change lightning site-directed mutagenesis kit (Strategene, USA) and the mutational primer 5’-GACTTTTCCTCTAGCAAGAGGATGATGCCCCAGTACCAGAAGGAGC-3’, as previously described (Martin *et al*., 2013). All constructs were sequenced at The Australian Genome Research Facility, located at The University of Queensland.

### Experimental model and subject details

**Mice** *DDHD2^-/-^* mice generated in a C57BL/6 background using standard gene targeting techniques (Inloes *et al*., 2014) were sourced from the Scripps Research Institute in the United States. The animals were maintained on a 12 h/12 h light/dark (LD) cycle at between 21-22 °C and housed in duos with access to standard mouse chow (in Dresden: Ssniff R/M-H; catalogue # V1534 and in Brisbane: Specialty Feeds, catalogue # SF00-100) and *ad libitum* autoclaved water.

*STXBP1*^+/-^ mice were generated as described previously (Chen *et al*., 2020) and housed in an Association for Assessment and Accreditation of Laboratory Animal Care International-certified animal facility at Baylor College of Medicine on a 14 h/10 h LD cycle. All procedures to maintain and use mice were performed in strict accordance with the recommendations in the Guide for the Care and Use of Laboratory Animals of the National Institutes of Health and were approved by the Institutional Animal Care and Use Committee at Baylor College of Medicine.

### Behavioural Experiments

#### Instrumental conditioning test

##### Animals

All behavioural and lipidomics experiments were performed using a cohort of age- and sex-matched *DDHD2^-/-^* and their *DDHD2^+/+^* littermates (3 and 12 months old, C57BL/6 background), while all other experiments were carried out using neurons from *DDHD2^-/-^* and C57BL/6 WT animals (Inloes *et al*., 2014) housed in 12 h/12 h LD environment with diet restricted to 85 % of their free-feeding weight. Every effort was made to minimize the number of animals used and any suffering. A total of 80 animals were used (20 mice per group) for behavioural experiments, while 6 animals were sacrificed per group, with brain samples collected for lipidomics analysis.

##### Apparatus

Instrumental conditioning training procedures occurred in sound and light resistant operant chambers (MED Associates) as previously described (Bertran-Gonzalez *et al*., 2013). In brief, the chambers were illuminated with a 3 W, 24 V house light. Each chamber contained a recessed feeding magazine in the centre of the chamber wall on one end. The magazine was connected to two pellet dispensers that could individually deliver grain and purified pellets (20 mg Dustless Precision Pellets; Bioserve Biotechnologies) into the magazine when activated. Either side of the magazine contained retractable levers. Med-PC (MED Associates) software was used to direct the insertion and retraction of the levers, illumination of the chamber, and delivery of the pellets. This software also recorded the number of lever presses, magazine entries, pellets delivered and duration of the experiment (30 minutes). Activity was monitored using a camera and D-View Cam Software (D-link Corporation; Taiwan).

##### Behavioural procedures

Behavioural training of the mice was carried out as previously described (Bertran-Gonzalez *et al*., 2013). In brief, the mice were trained to press a lever (left or right) to obtain a food reward (20 mg dustless precision pellets; Bioserve Biotechnologies, 3.35 kcal/g). A total of 10 mice were trained per session and two conditions were used: instrumental and control. The instrumental animals were presented with a lever and a reward contingent with the number of lever presses, while the control animals were only exposed to the box but not presented with a lever and there was no reward delivery. Each session ran for 30 min or until the animals earned 20 rewards (whichever came first).

##### Magazine training

Each mouse (*DDHD2^+/+^* and *DDHD2^-/-^*) was allotted to a specific operant chamber, which was maintained throughout the experimental period. For magazine training during the first 3 days (days -1 to -3), the instrumental group animals received a maximum of 20 pellets per mouse for the 30 min period. The chamber was illuminated to indicate the onset of the session and extinguished at the expiration. Following the training, both levers were retracted, and the mouse allowed to freely explore the chamber.

##### Instrumental training

Instrumental mice were trained on a continouos reinforcement schedule (CRF) where the delivery of the rewarding outcome was contingent on the corresponding lever-pressing action (one reward delivered for each lever press) for first 3 days of instrumental training (days 1-3). Thereafter, a random ratio (RR) schedule was initiated to reduce the chance of obtaining a reward. The mice were trained on RR5 for 3 days (outcome delivery for each action was at a probability of 0.2; training days 4-6), then RR10 for 3 days (probability 0.1; training days 7-9), and lastly RR20 for five days (probability 0.05; training days 10-14). Control mice were not presented a lever, and hence, no reward was delivered. After the 17 days of training, the memory performance for each mouse was ascertained by computing the lever press rate (lever presses per minute).

##### Novel object location (NOL) test

The NOL paradigm is a variant of the novel object recognition (NOR) test (Lueptow, 2017) which affords assessment of spatial memory based on the ability of the animal to encode both the object’s feature as well as the spatial location information of the object (Massey *et al*, 2003). In this paradigm, all mice were habituated to the room as well as to two identical objects placed in opposite locations of a white box (30.5 × 30.5 cm) for 10 min. After 24 hours, one of the objects was moved to a novel location and the mice were allowed to explore the objects in the different location for 10 min while being videoed. The time that the mice spent exploring the NOL (discrimination index), defined as exploration with the nose, less than 1 cm from the object, was quantitatively assessed using software (Ethovision) (Blackmore *et al*., 2022).

##### Activity monitoring test

Using an automated activity monitoring device (Med Associates), we assessed open field locomotor performance, alongside explorative and vigilance activity. Briefly, each mouse was placed in the centre of an arena (40 × 40 × 30 cm) and allowed to freely explore in the presence of approximately 70-lux illumination throughout the testing period. Parameters including vertical time, resting time, jumping time, vertical counts, jump counts, and ambulatory distance, were recorded over a 30-minute trial period (Fan *et al*, 2023; Pandit *et al*, 2019).

##### Grip strength test

The grip strength for both forelimb and hind limb was assessed monthly (between 3 and 12 months of age) using a Digital Force Gauge (Ugo Basile, System; San Diego Instruments Inc., San Diego, CA). Each mouse was held close to the base of its tail and placed on the rectangular bar of the grip strength device, ensuring that all four paws made contact before gently pulling at a consistent angle, to automatically record the peak force in Newtons before the animal’s grip was broken. A total of 10 trials per mouse were carried out and the mean peak force of the 10 trials was used for subsequent analysis (Pandit *et al*., 2019).

##### Motor coordination test

Motor coordination was assessed monthly using an accelerating Rota-Rod device (Ugo Basile, Comerio, Italy) by determining the latency period from when the mouse is placed on the accelerating rotating rod device to the initial fall (the length of time the mouse lingered on the rolling rod). A total of three trials was carried out for each mouse, at an acceleration mode of 5 to 30 rpm over 60 s. For the three individual trials, the latency to the first fall was averaged and used as the mean latency, while a ceiling latency of 180 s was recorded if the mouse persisted beyond this period (Pandit *et al*., 2019).

### Tissue Collection

To determine the FFA response, approximately 4 hours following instrumental conditioning the mice were sacrificed under deep anaesthesia (intraperitoneal injection of 0.2 ml phenobarbitol (300 mg/ml)) and transcardially perfused with ice-cold phosphate-buffered saline (PBS) pH 7.4 to flush out blood and chill the brain prior to snap freezing in liquid N_2_. Tissue was then cryodissected from 5 different frozen brain regions (prefrontal cortex - PFC, ventral hippocampus – VH, dorsal hippocampus - DH, amygdala - AM, and cerebellum - CB) to minimize post-mortem and ischemic lipid metabolism (Wallis *et al*., 2021). The animals were swiftly sacrificed, and their brains were excised, carefully wrapped in aluminium foil and snap-frozen in liquid nitrogen (subsequently stored at -80 °C). Frozen brains were mounted on the block of a cryostat (NX70, Thermo Fisher Scientific) running at -20 °C, where 80-100 µm thick tissue slices were cut and transferred to a glass slides (Merck; S8902). Slides were then transferred to an Olympus SZ51 microscope next to the cryostat, and suitable brain areas were quickly dissected from each frozen slice and placed on dry ice in 2 ml tubes (Eppendorf). Each brain was split into 70 slices on average. Material was dissected from roughly 20 sequential brain slices in each tube, which represented a specific brain region. Until the lipid extraction, all samples were kept at -80 °C. STXBP1^+/-^ mouse tissues were treated in a similar way and the extraction operations were also carried out in frigid temperatures minimize ischemic and post-mortem lipid metabolism, and at no stage prior to this were brain samples allowed to thaw.

### Synthesis of FFAST Isotopic-Coded Differential Tags

From commercially available CH_3_I/CD_3_I/C_2_H_5_I and 3-hydroxymethyl-pyridine reagents, 3-hydroxymethyl-1-methylpyridium iodide (FFAST-124), 3-hydroxymethyl-1-methyl-d3-pyridium iodide (FFAST-127), and hydroxymethyl-1-ethylpyridium iodide (FFAST-138) tags were produced. The synthesis of FFAST derivatives was carried out as described previously (Koulman *et al*, 2009; Narayana *et al*., 2015). In brief, 200 mg of iodomethane, iodomethane-D3, and iodoethane were each mixed with 100 mg of 3-hydroxy-methyl-pyridine. Under gaseous N_2_, the resulting solution was heated to 90 °C and microwaved for 90 min at 300 W in a CEM Discover microwave reactor. The resultant solution was then dried after being rinsed with 100 % diethyl ether. Using 1H nuclear magnetic resonance (NMR) spectroscopy, the purity of all the derivatives was determined to be > 95 %.

### Free Fatty Acid (FFA) extraction and FFAST labelling of brain tissue

In the cold room, frozen brain tissue samples were homogenised for 5 min in 0.5 mL of HCl (0.1 M) using a tissue homogenizer ultrasonic processor (Vibra-Cell, Sonics Inc. USA). 200 mL homogenate was treated with 0.6 mL ice-cold chloroform and 0.4 mL ice-cold methanol:12 N HCl (96:4 v/v, supplemented with 2 mM AlCl_3_). After the mixture had been thoroughly vortexed, 0.2 mL ice-cold water was added, and tubes centrifuged at 4 °C for 2 min at 12000 x g in a refrigerated microfuge (Eppendorf 5415R). The upper phase was discarded, and the lower phase tube was dried in a vacuum concentrator (Genevac Ltd). Dried extracts were re-dissolved in 100 μL acetonitrile. The derivatization strategy was designed for molecules with free carboxylic acid groups and followed a previously published procedure (Narayana *et al*., 2015). Briefly, 100 μL of FFA extracts in acetonitrile were combined with 50 μL of 1,1-carbonyldiimidiazole (1 mg/mL in acetonitrile) and incubated at room temperature (RT) for 2 min. Following this, 50 μL of either FFA extracts tagged with FFAST-124 or FFAST-127 (50 mg/mL in acetonitrile, 5 % triethylamine) was added. The combinations were then mixed for 2 min before heating for 20 min at 50 °C in a water bath. Finally, 100 μL of each isotopically labelled sample was combined and dried in a vacuum concentrator. They were then redissolved in 200 μL of an internal standard solution (2.5 μM in acetonitrile) made by derivatizing the 19 FFA standards with the FFAST-138 label. The samples were then placed in an auto-sampler vial and analysed using liquid chromatography tandem mass spectrometry (LC-MS/MS). Prior to analysis, all samples were maintained at -20 °C.

### FFA LC-MS/MS Analysis

LC-MS/MS analysis was performed on a Shimadzu Nexera UHPLC equipped with an Poroshell 120 CS-C18, 2.1 × 100 mm column with 2.7 µm particle size, linked to an AB Sciex 5500QTRAP tandem mass spectrometer fitted with an ESI Turbo V source. Analyst ® 1.5.2 software (AB Sciex) was used for instrument control data collection while Multiquant software (AB Sciex) was used for the multiple reaction monitoring (MRM) data analysis. Chromatographic separations were performed on 1 µL sample injection volume using a gradient system consisting of solvent A (0.1 % formic acid (v/v)) and solvent B (100 % acetonitrile with 0.1 % formic acid (v/v)) was used to perform liquid chromatography (LC) at 0.450 mL/min heated to 60 °C). The gradient conditions consisted of an initial isocratic step of 15 % B for 1 min followed by a gradient to 100 % B over 9 min. The column was flushed at 100 % B for 2 min and then reduced back to 15 % to re-equilibrate for 2 min for a total run time of 14 min. The first 0.5 min of the LC run was switched to waste to remove any excess underivatized FFAST tags.

Mass spectrometric data acquisition was performed using positive mode ionisation in MRM mode (Table S1, S2, and S3). Ion source temperature was set at 400 °C, and ion spray voltage set to 5500 V. The source gases setting consisted of curtain gas, GS1 and GS2 set to 45, 40 and 50 psi respectively. The collision energy, declustering potential, and collision cell exit potentials were all set at 50, 100, and 13 V, respectively for all transitions.

### Cell Cultures

PC12 cells were cultured in Dulbecco’s modified Eagle’s media (DMEM) supplemented with glucose (4500 mg/L) and containing 30 mM NaHCO3, 5 % foetal bovine serum, 10 % horse serum, 100 units/ml penicillin/streptomycin, and 100 g/ml Kanamycin, and maintained as described previously (Han *et al*., 2009; Martin *et al*., 2013). STXBP1/2 DKO PC12 cells were created using CRISPR (Kasula *et al*.). Production of STXBP1^F115E^, STXBP1^Δ317-333^ and STXBP1^WT^ recombinant proteins were produced as previously described (Christie *et al*, 2012; Deak *et al*., 2009; Kasula *et al*., 2016; Martin *et al*., 2013).

### Immunofluorescence microscopy

PC12 and STXBP1 DKO PC12 cells were transfected with STXBP1^WT^, STXBP1^Δ317-333^, and STXBP1^F115E^ constructs and immunolabelling was carried out as previously described (Malintan *et al*., 2009; Martin *et al*., 2013). Following permeabilization of cells using 0.1 % Triton X-100, cells were imaged using a Zeiss LSM510 confocal microscope.

### Secretagogue stimulation of PC12 and DKO-PC12 cells

Cells transfected with STXBP1^WT^, STXBP1^Δ317-333^, STXBP1^F115E^, or STXBP1^C180Y^ constructs tagged with emGFP were washed with Buffer A (145 mM NaCl, 1.2 mM Na_2_HPO_4_, 20 mM HEPES-NaOH, 2 mM CaCl_2_, 10 mM glucose, pH 7.4) and stimulated with either 60 mM KCl or 2 mM KCl and incubated for 15 min at 37 °C. Cells were collected, and FFAs were extracted using chloroform-methanol as described above. The FFAs were redissolved in 100 μL of acetonitrile.

### Immunoprecipitation (Pull-Down assay)

PC12 and STXBP1 DKO PC12 cells were maintained in DMEM (containing sodium pyruvate; Thermo Fisher Scientific), foetal bovine serum (7.5 %, Gibco) and horse serum (7.5 %, Gibco), and 0.5 % GlutaMax (Thermo Fisher Scientific) at 37 °C an 5 % CO_2_. Cells were transfected using Lipofectamine®LTX with Plus Reagent (Thermo Fisher Scientific) as per the manufacturers’ instructions with 2 ug of Munc18-1-GFP and DDHD2-GFP; respectively. Non-GFP transfected wild type cell lines where prepared in parallel as a process control. 48 h post-transfection, PC12 cells were incubated for 5 min in isotonic buffer A (145 mM NaCl, 5 mM KCl, 1.2 mM Na2HPO4, 10 mM D-glucose, and 20 mM Hepes, pH 7.4) or stimulated with 2 mM BaCl_2_ buffer A for 5 min at 37 °C. Cells were homogenised in ice cold lysis buffer (10 mM Tris/HCl pH 7.5; 150 mM NaCl; 0.5 mM EDTA; 0.5 % NP-40, and EDTA-free protease inhibitor cocktails; Roche) for 15 min on ice and the cell lysate was centrifuged at 13000 RPM for 15 min, the supernatant was transferred to a new tube containing 25 µl of anti-GFP magnetic beads and equilibrated twice with washing buffer (10 mM Tris/HCl pH 7.5; 150 mM NaCl; 0.5 mM EDTA). During the tumble end-over-end rotation for 1 h at 4 °C, GFP-Munc18-1 or DDHD2-GFP is captured by the anti-GFP magnetic beads. The magnetic beads were then washed three times in washing buffer and magnetically separate until supernatant became clear and all unbounded protein completely washed out. Bead-bound samples for Western Blot analysis were resuspended in 100 μl 2X SDS-sample buffer (120 mM Tris/HCl pH 6.8; 20 % glycerol; 4 % SDS, 0.04 % bromophenol blue; 10 % β-mercaptoethanol), whereas sample for LC-MS/MS proteomics analysis were suspended in SDS-PAGE buffer with the following modifications buffer (50 mM Tris-HCl pH 6.8, 10 mM dithiothreitol, 2 % w/v sodium dodecyl sulphate, 10 % v/v glycerol). Samples were then and boiled for 10 min at 95 °C to dissociate the formed immunocomplexes from the beads and then magnetically separated to remove to beads from the protein suspensions. Samples were stored at - 20^0^C until further analysis.

### Immunocytochemistry

PC12 cells and STXBP1 DKO PC12 cells were stimulated with 2 mM of BaCl_2_ buffer A for 5 min and fixed in 4 % paraformaldehyde for 30 min at RT. Cells were washed in PBS containing 0.2 % BSA and permeabilized with 0.1 % Triton X100 for 4 min. After permeabilization, non-specific binding sites were blocked with 1 % BSA in PBS for 30 min at RT. Primary and secondary antibodies were applied in blocking buffer (1 % BSA in PBS) for overnight (4 °C) and 1 h (RT); respectively. The following primary antibodies were used: anti-DDHD2 (1:50), anti-Munc18-1 mouse monoclonal antibody (1:50) and anti-syntaxin mouse monoclonal antibody (1:50).Secondary antibodies: Alexa Fluor® 488 conjugated anti-mouse IgG, Alexa Fluor® 546 conjugated anti-guinea pig IgG, Alexa Fluor® 546 conjugated anti-rabbit IgG and Alexa Fluor® 647 conjugated anti-rabbit IgG, were diluted 1:1000 in blocking buffer.

### Imaging and Image Analysis

Confocal images were acquired with a spinning-disk confocal system consisting of an Axio Observer Z1 equipped with a CSU-W1 spinning-disk head, ORCA-Flash4.0 v2 sCMOS camera, 63x 1.3 NA C-Apo objective. Image acquisition was performed using SlideBook 6.0. Sections of each slide were acquired using a Z-stack with a step of 100 nm. The exposure time for each channel was kept constant across all imaging sessions. Images were deconvolved on Huygens deconvolution software 21.10 (Scientific Volume Imaging). Both channels are thresholded using Costes method before calculating the Pearson’s coefficient. For individual cells for rescues, the Costes method was applied only on the region of interest (ROI) corresponding to the cell (based on GFP-Munc181 signal).

### Western blot

Briefly, the pulled down proteins were electrophoresed on 4-20 % precast polyacrylamide gel (BIO-RAD) for 1 h with 100 fixed voltage and transferred to PVDF membranes using wet method for 90 min with100 fixed voltage. Membranes were then blocked with intercept blocking buffer (LI-COR) for 1 h at RT. Membranes were then incubated with DDHD2 rabbit polyclonal primary antibody (Proteintech #25203-AP; 1:500) and anti-Munc18-1 mouse monoclonal antibody (BD Biosciences #610337; 1:1000) overnight at 4 °C, followed by 1 h incubation with the anti–mouse IgG and anti-rabbit HRP-secondary IgG antibodies (Cell Signalling). Then, proteins were then visualized by ECL detection solution using LI-COR system.

### Quantitative real time PCR

PC12, and STXBP1/2 DKO PC12 cells were collected for RNA isolation. RNA was isolated using RNeasy mini kit (Qiagen). 1 µg of RNA was reverse transcribed to cDNA using High-capacity reverse transcription kit (Applied Biosystems). RT-PCR was carried out using the following primers: rat DDHD1 forward 5′-CATCGATGGAAAAGACGCTGT-3′ and reverse 5′-CCCACTGCTGGAGGCTTTAG-3′, rat DDHD2 forward 5′-CATCGATGGAAAAGACGCTGT-3′ and reverse 5′-CCCACTGCTGGAGGCTTTAG-3′, rat beta actin forward 5′-CCCGCGAGTACAACCTTCTTG-3′ and reverse 5′-GTCATCCATGGCGAACTGGTG-3′. Using PowerUpnTM SYBRTM Green Master Mix (Applied Biosystems) and using the following cycling conditions (denaturation at 95 °C for 15 s, annealing at 60 °C for 30 s, and extension at 72 °C for 30 s) for 40 cycles. The mRNA levels were normalized to β-actin mRNA levels and estimation as delta-delta cycle threshold (DDCT) was calculated as described previously (Rao *et al*, 2013). All Real-Time Quantitative Reverse Transcription PCR (qRT-PCR) reactions were carried out in triplicate in each biological replicate and each product size was confirmed with the agarose electrophoresis.

### NPY-hPLAP release assay

The neuropeptide-Y-human placental alkaline phosphatase (NPY-hPLAP) release assay was carried out according to a previously described procedure (Martin *et al*., 2013). PC12 cells and STXBP1 DKO PC12 cells were co-transfected for 72 h with NPY–hPLAP and the corresponding STXBP1 plasmids. Cells were washed and incubated for 15 min at 37 °C with PSS buffer (5.6 mM KCl, 145 mM NaCl, 0.5 mM MgCl_2_, 2.2 mM CaCl_2_, 15 mM Hepes-NaOH, 5.6 mM glucose, pH 7.4) or depolarized with high K^+^ - PSS buffer (2.2 mM CaCl_2_, 70 mM KCl, 0.5 mM MgCl_2_, 81 mM NaCl, 15 mM Hepes–NaOH, 5.6 mM glucose, pH 7.4) and incubated for 15 min at 37 °C. To measure NPY–hPLAP release, supernatant was collected, and cells were lysed with 0.2 % Triton X-100. Using the Phospha-Light^TM^ chemiluminescent reporter gene assay system (Applied Biosystems), the NPY-hPLAP release and total were determined.

### ALPHAScreen protein interaction assay

Amplified Luminescent Proximity Homogeneous Assay Screen (ALPHAScreen) was carried out utilising the cMyc detection kit and Proxiplate-384 Plus plates (PerkinElmer) as previously described (Martin *et al*., 2013; Sierecki *et al*., 2013). Each protein pair (one tagged with C-terminal GFP and the other with C-terminal Cherry-cMyc) was co-expressed in 10 μl of *Leishmania tarentolae* extract (LTE) for 3 h at 27 °C using 20 and 40 nM of DNA template, respectively. LTE lysate co-expressing the proteins of interest was diluted in buffer A (25 mM HEPES, 50 mM NaCl). Each sample was subjected to a four-fold serial dilution. Anti-cMyc coated Acceptor Beads were aliquoted into each well in buffer B (25 mM HEPES, 50 mM NaCl, 0.001 % NP40, 0.001 % casein) for the experiment. 2 μl of diluted sample and 2 μl of biotin labelled GFP-Nanotrap were then added to buffer A. The plate was incubated at RT for 45 min. 2 μl (0.4 μg) of Streptavidin-coated Donor Beads were added, diluted in buffer A, and incubated in the dark for 45 min at RT. Using an Envision Multilabel Plate Reader (PerkinElmer), the ALPHAScreen signal was captured using the manufacturer’s suggested settings (excitation: 680/30 nm for 0.18 s, emission: 570/100 nm after 37 ms). A positive interaction is indicated by the formation of a bell-shaped curve, whereas a flat line indicates a lack of contact between the proteins. Each protein pair was measured a minimum of three times using different plates each time.

### Proteomics analysis: Un-targeted protein Identification using High Resolution Tandan Mass spectrometry

Sample processing of immuno-precipitates from GFP pull-downs and wild type process controls in modified SDS-PAGE buffer consisted of protein denaturation and disulfide bonds reduction by heating to 60 °C for 30 min in a heating block. Cysteine residues were then alkylated to prevent disulfide bond reformation using 50 mM iodoacetamide for 45 min at RT in the dark. Proteins were then precipitated through the addition of 10 volumes of ice-cold 1:1 methanol/isopropanol followed by overnight incubation at -20 °C. Precipitates were then pelleted by centrifugation at 14,000 *g* for 15 min in a benchtop microfuge. The supernatant was removed, and the pellet was washed with the addition of 1 mL of ice-cold 1:1 methanol/isopropanol followed by a second round of centrifugation as previously described. The pellet was resuspended in 20 uL of 50 mM ammonium bicarbonate containing 2 µg of proteomics grade trypsin (Sigma Aldrich). Proteolytic digestion was performed overnight at 37 °C in a shaking incubator set to 200 rotations/minute.

Unknown protein identification was performed by injecting 9 µL of the peptide digest onto a 5600 TripleTOF mass spectrometry (AB Sciex) with a microflow LC (Eksigent). Chromatographic conditions consisted of first injecting the sample onto a 0.3 × 10 mm C18 micro trapping column (Phenomenex) under an isocratic flow of 0.1 mM formic acid at 5 µL/min. After 10 min the trapping column was then switched in-line with the separation column (CL120 0.3 × 150 mm C18 with 3 µm particle size, Eksigent). Gradient mobile phases consisted of A) 0.1 mM formic acid and B) acetonitrile containing 0.1 % formic acid, at a flow rate of 5 µL/min. Chromatographic separations started at 5 % B) at progressing to 32 % B) at 68 min, 40 % B) at 72 min, and 95 % B) at 76 min, plateauing until 79 min, before dropping to 3 % B) at 80 min followed by aqueous column re-equilibration for 7 min. MS acquisitions were performed using positive electrospray ionization using information-dependent data acquisition (IDA) mode. Source conditions consisted of curtain, GS1, and GS2, gases set to 30, 30, and 20 psi respectively, source temperature at 250 °C, and ion spray and declustering potentials set to 5500 and 100 V respectively. IDA data acquisition was performed using an initial MS survey scan for 250 ms. From this, the top 30 most abundant ions with an intensity greater than 100 cps, a mass exceeding 350 Da and a charge state between 2-5 were selected for MS/MS collision-induced dissociation (CID) fragmentation. CID collision energy for each of these ions was calculated using the rolling collision energy algorithm, scaling to ion mass. MS-MS spectra were acquired between 100 and 2000 *m/z* for 55 ms.

Data and analysis and protein identification was performed using the Protein Pilot software (ABSciex) searched against the combined SwissProt and TrEMBL *Rattus norvegicus* proteomes (31,557 total entries) obtained from the Uniprot online repository (Consortium, 2021). Protein pilot parameters consisted of performing a thorough search, cystine modification with iodoacetamide, and digestion with trypsin, using an ID focus on biological modification with the default paragon method settings for modification frequency. Protein identifications were screened statistically using false discover rate (FDR) using a 1 % global threshold for positive identification. Positively identified proteins were then further graded according to the number of identified peptides after 99 % confidence screening. Proteins positively identified in the wild-type process controls were considered to present due to non-specific interactions and removed from the list of proteins identified from the pulldown assays. Identified proteins with 2 or more peptides identified then underwent interaction and gene ontology enrichment analysis using the Search Tool for Retrieval of Interacting Genes/Proteins (STRING) Database online resource (Szklarczyk *et al*., 2021).

### Proteomics Analysis: Targeted quantification of DDHD2 and STXBP1 using Multiple Reaction Monitoring

A bespoke targeted multiple reaction monitoring (MRM) LC-MS/MS assay was employed for quantification of *Rattus norvegicus* DDHD2 and STXBP1. Assay design consisted of obtaining protein sequence information for rat isoforms for DDHD2 (D3ZJ91) and STXB1 (P61765 and P61765-2) from the Uniprot database. Modelling of *in silico* tryptic digestion of these proteins, calculation of *b* and *y* fragment ion masses and theoretical optimal collision energies for each transition using an AB Sciex QTRAP mass spectrometer was determined using the Skyline software platform (MacCross Laboratory) Putative peptides were then screened for specificity to the target protein using blastp function of the NCBI online resource. Finally, MRM transitions for proteotyptic peptides were screened for visibility using the LC-MS/MS methodology described below using tryptic digests of whole cell protein extracts from rat PC12 cells. The LC-MS/MS specificity of peptides yielding an unambiguous chromatographic peak was confirmed by matching MS2 fragmentation spectra obtained from enhanced product ion scans of PC12 lysates to at least three of the modelled fragment ion masses for each peptide obtained from Skyline.

MRM analysis was performed on an AB Sciex 5500 QTRAP mass spectrometer with a Shimadzu Scientific Nexera series liquid chromatography system. Chromatographic separations were performed using a 3 µL injections volume of the tryptically digest immuno-precipitates used for high resolution protein identification onto a Aeris peptide XB-C18 2.1 × 100 mm column with a 2.5 µm particle size. Mobile phases consisted of A) 0.1% v/v formic acid and B) acetonitrile modified with 0.1% formic acid at a flow rate of 0.5 mL/min. Separations were performed over 20 min using gradient conditions consisting of an initial isocratic step at 5% B for three min followed by a shallow gradient from 5% B to 50% B at 15 min followed by column washing by increasing to 100% B at 17 min, holding for 3 min and a final column re-equilibration at 5% B for 2 min. Mass Spectrometric acquisitions were performed using positive mode electrospray ionisation. Ion sources gases consisted of curtain gas, GS1 and GS2 applied at 30, 50 and 60 psi respectively. Source temperature was set to 550^0^C. Ion path potentials consisted of ion spray (5500 V), declustering potential (80 V), entrance potential (10 V) and exit potential (9). Dwell time for each transition was 20 ms for a total cycle time of 800 ms. Presursor *m/z,* product *m/z* and collision energy for each transition is displayed in the table S4.

### Electron microscopy

Electron microscopy (EM) analysis was performed on cultured embryonic day 16 (E16) C57BL6 and DDHD2^-/-^ hippocampal neurons grown on glass bottom dishes (CellVis, Cat # D29-10-1.5-N) at 100,000 neurons per 78.54 mm^2^ confluency to ensure mature and functional synaptic connections. Neurons were transfected with HRP-KDEL (Li *et al*, 2010) plasmid at 14 days in vitro (DIV14). On DIV19-20, neurons were fixed with 2 % paraformaldehyde, 1.5 % glutaraldehyde (Electron Microscopy Sciences, Cat #16210) in 0.1 M sodium cacodylate (Sigma-Aldrich, Cat #C0250) buffer pH 7.4 for 20 min at RT, washed three time for 3 min with 0.1 M sodium cacodylate buffer, and processed for DAB cytochemistry using standard protocols. Samples were contrasted with 1 % osmium tetroxide and 2 % uranyl acetate before dehydration and embedded in LX-112 resin using BioWave tissue processing system (Pelco) as previously described (Rigoni *et al*, 2005). Thin sections (80-90 nm) were cut using an ultramicrotome (Leica Biosystems, UC6FCS) and were imaged with a transmission electron microscope (JEOL USA, Inc. model 1101) equipped with cooled charge-coupled device camera (Olympus; Morada CCD Camera). Images were acquired when HRP-precipitate was observed. For the quantification of the percentages (%) of total HRP-signal, the localization of the HRP-tagged KEDL was categorized.

### Quantitative Data Processing and Visualization

Multiquant® 3.03 (AB SCIEX) was used to quantitatively analyse FFA, DDHD2 and STXBP1, with the MQ4 peak picking / integration algorithm used to manually quantify the LC-MS/MS peak area for each analyte. Quantification of FFAs was performed using a previously established calibration curve for that species, the abundance of each FFA (FFAST-124 or FFAST-127 labelled) was calculated relative to the FFAST-138 labelled internal standard for that species. Six serial dilutions of the FFAST-124 or FFAST-127 labelled FFA calibrators (0.05 ng/ml to 25 ng/ml in acetonitrile) were coupled with 2.5 ng/ml of the corresponding FFAST-138 labelled internal standard to obtain triplicate calibration curves. The ratio between analyte and internal standard peak area was plotted as a standard curve for each FFA. Linear regression without weighting was used to produce calibration slopes (R^2^ > 0.99). FFA quantification was carried out on experimental data using bespoke Python (python.org) programming. The concentration (ng/ml) of the analyte in the injected sample was calculated by dividing the ratio of the analyte (FFAST-124 or FFAST-127) peak area to the internal standard (FFAST-138) peak area by the calibration slope for that analyte, which was then converted to the molar concentration (pmol/μl) by dividing by the molecular weight of the labelled analyte. The concentration was then normalised to pmol/mg tissue using the weight of the tissue sample and the extraction volume.

Relative quantification techniques were used to assess DDHS2 and STXBP1 LC-MS/MS responses between treatment conditions. In brief this consisted of summing the peak areas for specific peptides across all treatment conditions to determine the relative response of that peptide for each condition. The response for each peptide of a protein were then summed and the aggregate used a measure of protein abundance. Statistical analysis and visualisation of aggregate protein data was performed using Prism® 9.0 (Graphpad).

To visualize and assess the significance of changes in analyte concentration across brain regions and in response to experimental conditions, lipid data were further analysed using custom-written Python (python.org) scripts variously utilizing Matplotlib (matplotlib.org), Pandas (pandas.pydata.org), Numpy (numpy.org), Seaborn (seaborn.pydata.org) and Scipy (scipy.org) modules. For each analyte the measurements for all the animals were processed to remove outliers by median filtering and generate the mean and standard error of the mean (SEM). Stacked barplots of these data were generated using Pandas and Matplotlib. Heatmaps were generated using Pandas and Seaborn. For heatmaps the significance of the fold-change for each pixel was determined by Student’s two-tailed t test using scipy.stats.ttest_ind. Scatterplots were generated using Pandas and Matplotlib.

All animal behavioural data was analysed and visualized using Prism® 9.0 (Graphpad).

### Main Text Statements

## Acknowledgements

This work was supported by Australian Research Council’s (ARC) Discovery Early Career Research Award (DECRA) to M.J. (DE190100565). F.A.M. is a National Health and Medical Research Council (NHMRC) Senior Research Fellow (GNT1155794). This work was also supported by an NHMRC Ideas Grant (2010901) awarded to F.A.M. and T.P.W. and a Clem Jones Centre for Ageing Dementia (CJCADR) Flagship Grant awarded to F.A.M. The Australian Government Research Training Program (RTP) Scholarship awarded to I.O.A., S.H., and B.M., by The University of Queensland, and a top-up scholarship courtesy of the Queensland Brain Institute (QBI) also supported this research. M.X. is a Caroline DeLuca Scholar. We thank Alex McCann for the critical reading of this manuscript. Figure 1a and extended Figure 1d were created using BioRender.

## Author Contributions

F.A.M., T.P.W., M.J. and I.O.A. conceived the study. I.O.A. performed all behavioural experiments with inputs from D.G.B., J.B-G., and M.J. B.G.V. and I.O.A. implemented the FFA and LC-MS/MS protocol and performed the lipidomic data acquisition as part of a PhD program under the supervision of F.A.M., T.P.W. and M.J. with input from B.G.V. B.M performed proteomics analysis. M.J. performed the electron microscopy analysis. A.A.M. and M.X provided the STXBP1^+/-^ mouse brains. S.H.S performed tissue culture, transfection, immunoprecipitation, western blotting, and immunocytochemistry staining and microscopy. R.S.G. performed tissue culture, transfection, and release assays. E.S. and Y.G. performed the ALPHAScreen assay. T.P.W. and I.O.A. performed the lipidomic data analyses. I.O.A. wrote the manuscript with input and revision from F.A.M., T.P.W., M.J., R.S.G., M.X., J.B-G., and B.C.

## Data availability

The data generated during this study, may be downloaded from the publicly accessible The University of Queensland Data Collection: Source data are provided with this paper.

## Code availability

The Python scripts for quantitative and multivariate data analysis and visualization are available upon request. Requests for software should be addressed to (F.A.M.).

## Extended Data Figure Legends

**Extended Data Figure 1.**
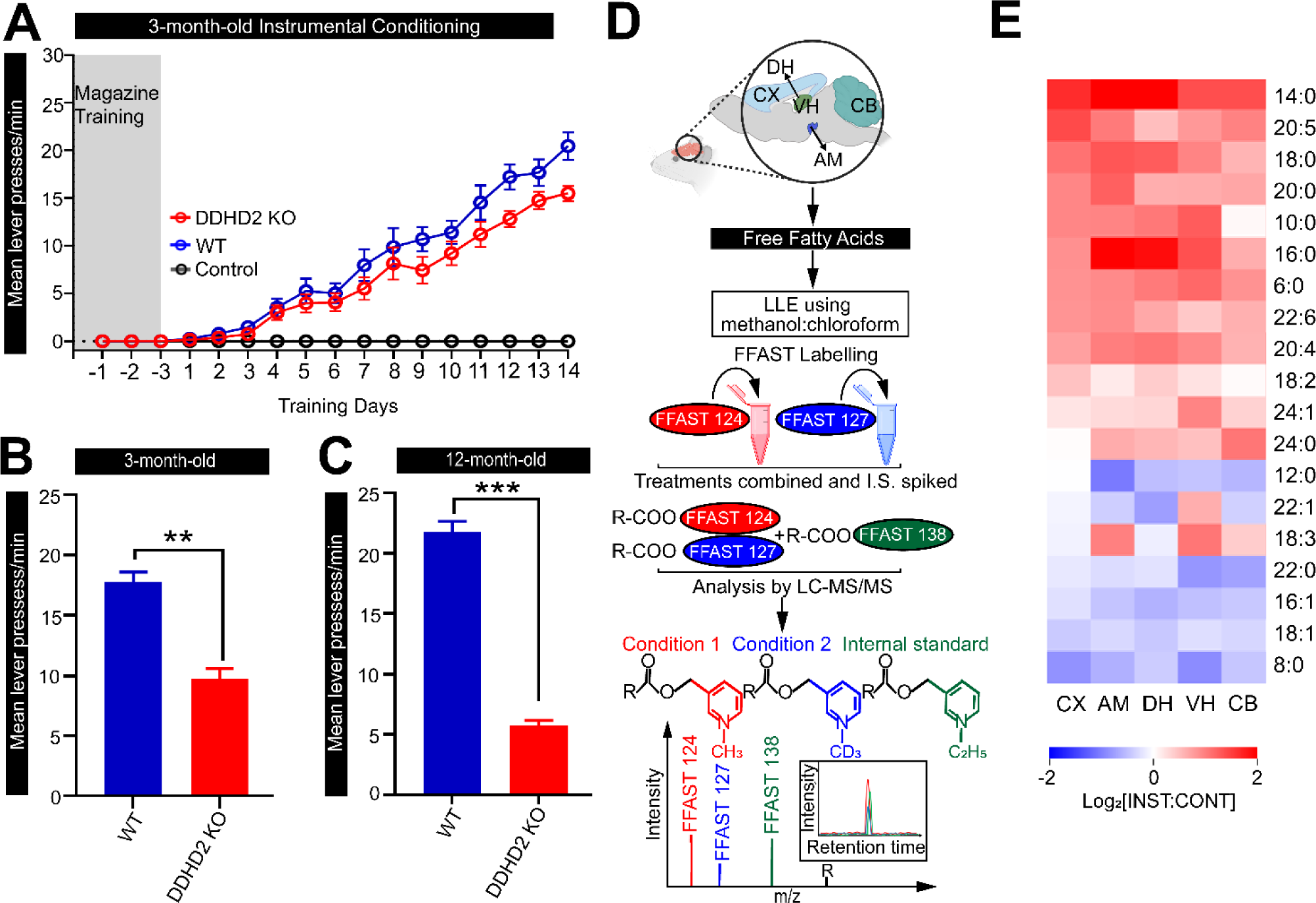
Long-term memory performance and FFA response in instrumentally conditioned mice. **A** Line graph showing mean lever presses per minute in 3mo WT vs KO mice (n=20; from two cohorts of animals). Bar graph showing behavioural response to instrumental conditioning in *DDHD2^+/+^* vs *DDHD2^-/-^* mice at **B** 3mo, and **C** 12mo. **D** Schematic workflow for Free Fatty Acid (FFA) extraction from different brain regions including cortex (CX), ventral hippocampus (VH), amygdala (AM), dorsal hippocampus (DH) and cerebellum (CB), and subsequent derivatization using FFAST, and LC-MS/MS analysis. For each brain region, FFAs were extracted from the two different experimental conditions (instrumental or context control mice) and labelled separately at the carboxy-terminus using FFAST for subsequent multiplexing LC-MS/MS analysis, together with FFAST-labelled internal standards (as per Narayana et al. 2015 (Narayana *et al*., 2015)). This workflow was replicated 6 times, to establish FFA abundance in each of the 6 mice used for each experimental condition. **E** Ordered heatmap showing the profile of 19 FFAs obtained by FFAST LC-MS/MS across 6 brains of 12-month-old control vs instrumental mice, with analytes shown by acyl chain composition. The significance of the difference between each group as determined by 1-way ANOVA (n=20) is indicated by asterisks ** p<0.01, *** p<0.001. Error bars represent the cumulative standard error of the mean (SEM) for all groups and parameters.

**Extended Data Figure. 2.**
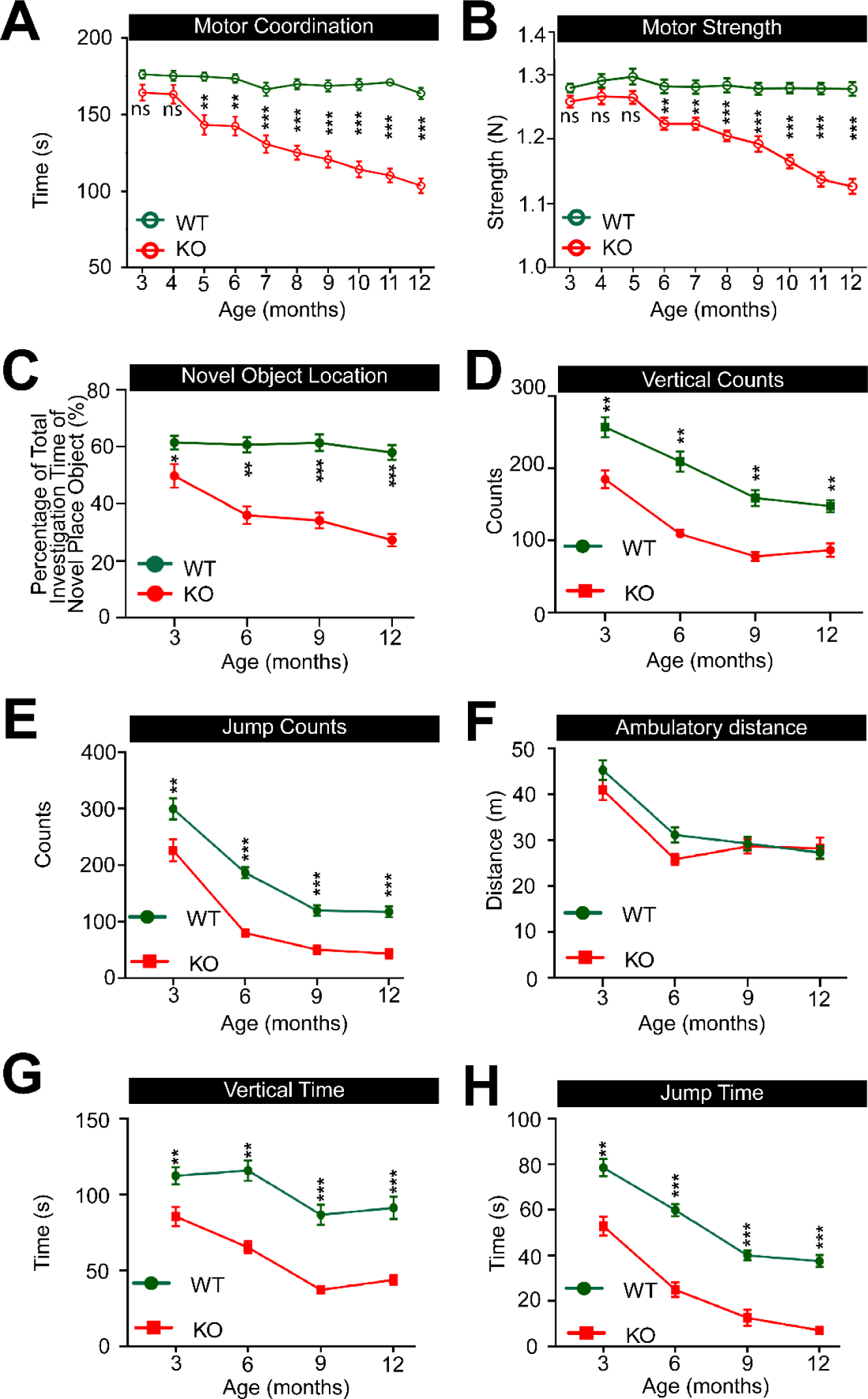
Longitudinal assessment of motor function, spatial memory, and explorative behaviors in *DDHD2^+/+^* vs *DDHD2^-/-^* mice. **A** Line graphs showing monthly recordings of **a** motor coordination, assessed using a rotarod device by determining the latency period from when the mouse is placed on the accelerating rotating rod device to the initial fall (s), and **B** motor strength (N), assessed using a grip strength device. **C** Longitudinal assessment of spatial memory performance in mice, using the Novel Object Location (NOL) paradigm, represented as a line graph. Longitudinal monitoring of mouse activity with line graphs showing **D** vertical counts, **E** jump counts, **F** ambulatory distance (m)**, G,** vertical time (s), and **H** jump time (s). The significance of the difference between each group as determined by 1-way ANOVA (n=20) is indicated by asterisks * p<0.05, ** p<0.01, *** p<0.001, ns = not significant. Error bars represent the cumulative standard error of the mean (SEM) for all groups and parameters.

**Extended Data Figure. 3.**
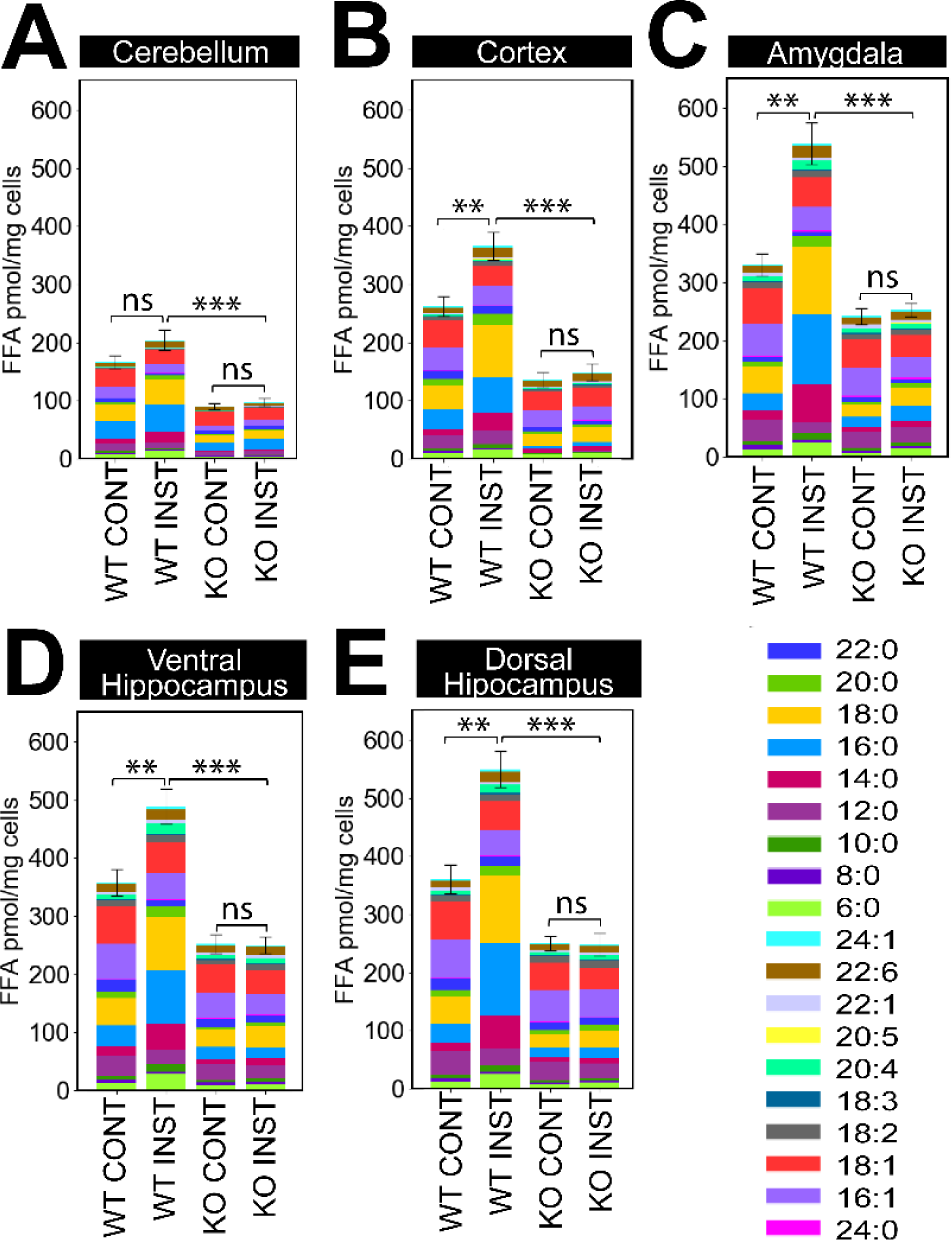
Total FFA levels across different brain regions of 12- month-old instrumentally conditioned *DDHD2^+/+^* vs *DDHD2^-/-^* mice. Bar plots show the profile of 19 FFAs obtained by FFAST LC-MS/MS across brains (n=6) of control vs instrumental mice, with analytes shown by acyl chain composition. The significance of the change in FFA abundance between the different experimental conditions as determined by one-way ANOVA with Holm-Sidak *post-hoc* correction is indicated by asterisks * p<0.05, ** p<0.01, *** p<0.001, ns = not significant. Error bars represent the cumulative standard error of the mean (SEM) for all groups and parameters. **A**, CB; Cerebellum, **B**, CX; Cortex, **C**, AM; Amygdala, **D**, VH; Ventral hippocampus, and **E**, DH; Dorsal hippocampus,

**Extended Data Figure. 4.**
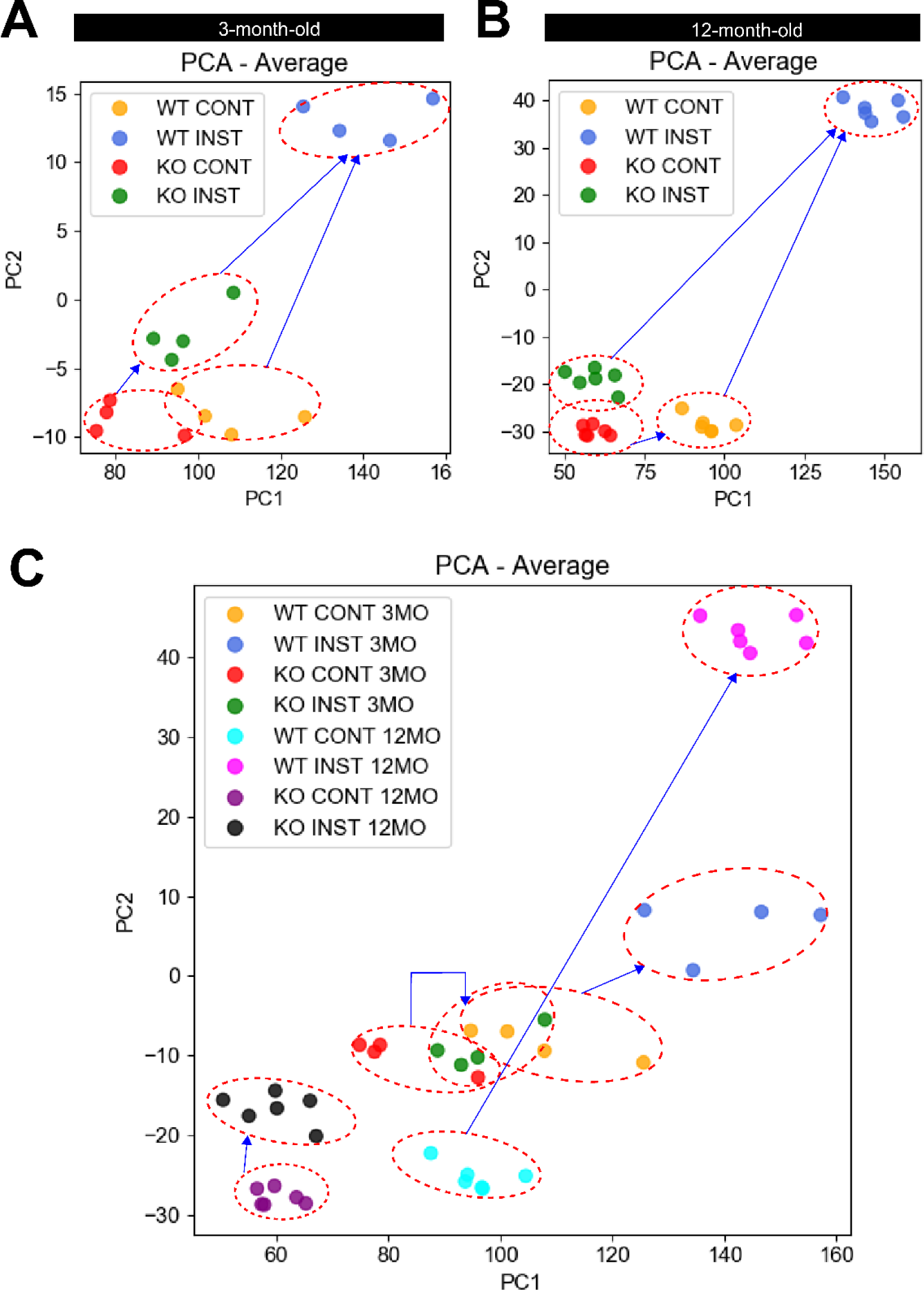
Average FFA response to instrumental conditioning in *DDHD2^+/+^* vs *DDHD2^-/-^* mice. The FFA profiles from all experimental animals was dimensionally reduced using PCA and clustered according to their similarity with a red dashed outline showing a clear distinction between the different groups (control WT mice, instrumental WT mice, control KO mice and instrumental KO mice) in order to show the average FFA response for each mouse. Each dot represents the normalized mean concentrations of the analytes observed across 6 animals for each condition. PCA analysis is shown for **A** 3mo mice, **B** 12 mo mice, and **C** 3mo and 12mo mice combined.

**Extended Data Figure 5.**
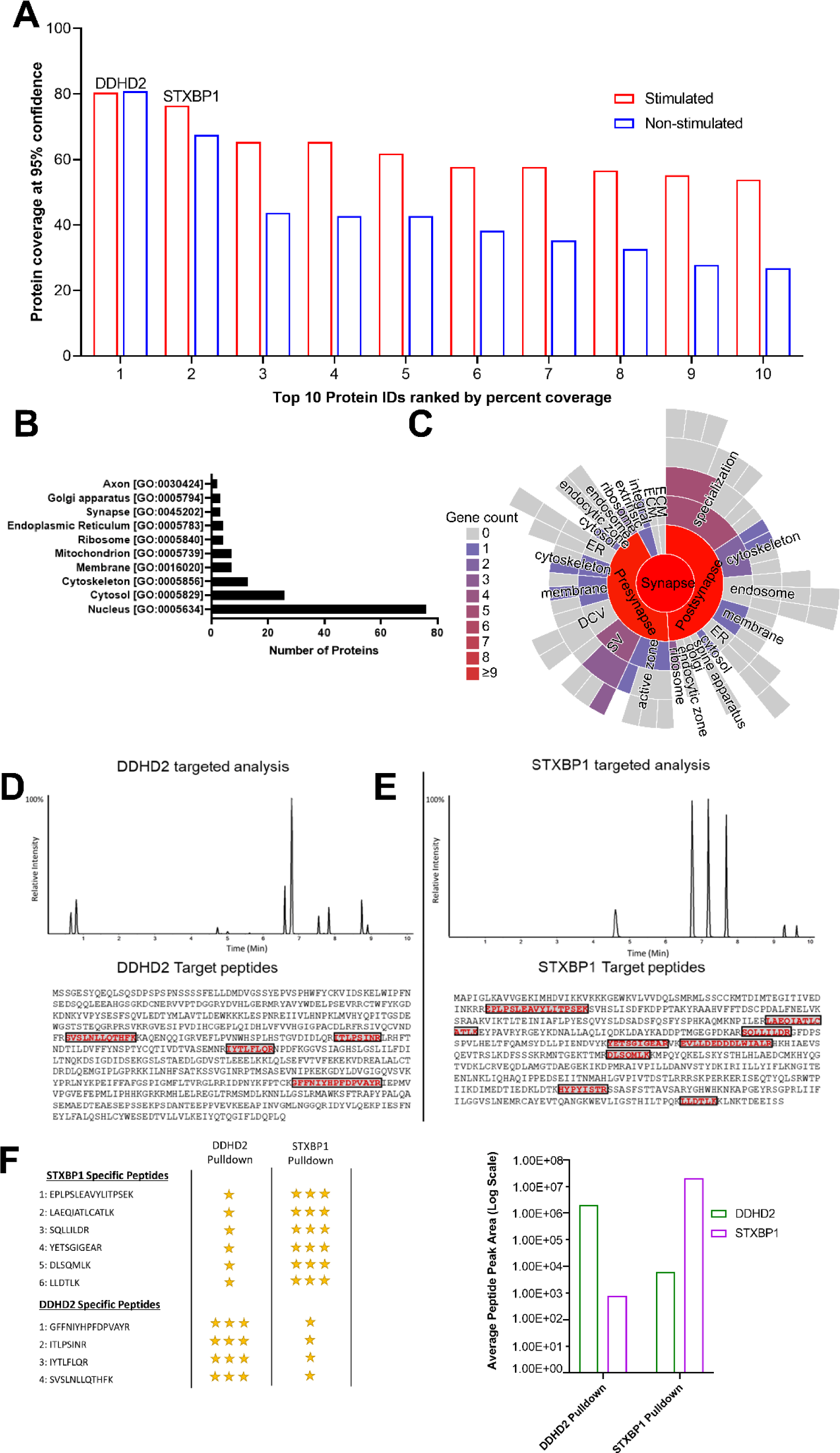
STXBP1 interacts with DDHD2. Untargeted high-resolution tandem mass spectrometry (HRMS) analysis of peptide digests from a DDHD2 pull down identified 34 proteins with two or more identified peptides at 1% local false discovery rate (FDR) in resting state unstimulated PC12 cells and 214 proteins in potassium excited cells. **A** Bar plot showing the top 10 proteins co-precipitating with DDHD2 from PC12 cell lysates identified at 1% FDR ranked by identified peptides sequence coverage at 95% confidence.. Using sequence coverage as an approximation of abundance Syntaxin binding protein 1 (STXBP1 the primary protein co-precipitating with DDHD2 in resting state and under potassium stimulation conditions. Cellular compartment gene ontology analysis of the identified proteins in potassium stimulated PC12 cells using uniprot database **B** shows that while the proteins co-precipitating with DDHD2 are primarily associated with the nucleus and cytosol, proteins from the synapse are detecting indicating that DDHD2 may play a role in synaptic function. Further cellular compartment gene ontology analysis specific for synaptic proteins using the SynGO database **C** detected membrane and cytoskeletal proteins associated with both the pre and post synapse. Interestingly proteins, such as STXBP1, associated with synaptic vesicles (SV) and the active zone in the pre-synapse were evident confirming an interaction between DDHD2 and neuronal exocytotic mechanisms. To confirm the interaction between DDHD2 and STXBP1 an orthogonal multiple reaction monitoring (MRM) assay using triple quadrupole tandem mass spectrometry was established, specifically targeting four proteotypic DDHD2 peptides **D** and six proteotypic STXBP1 peptides **E**. The qualitative abundance for each peptide in immunoprecipitation assays targeting DDHD2 and STXBP1 in PC12 cell lysates are displayed in panel **F** confirming the presence for both proteins in both pulldowns.

**Extended Data Figure 6.**
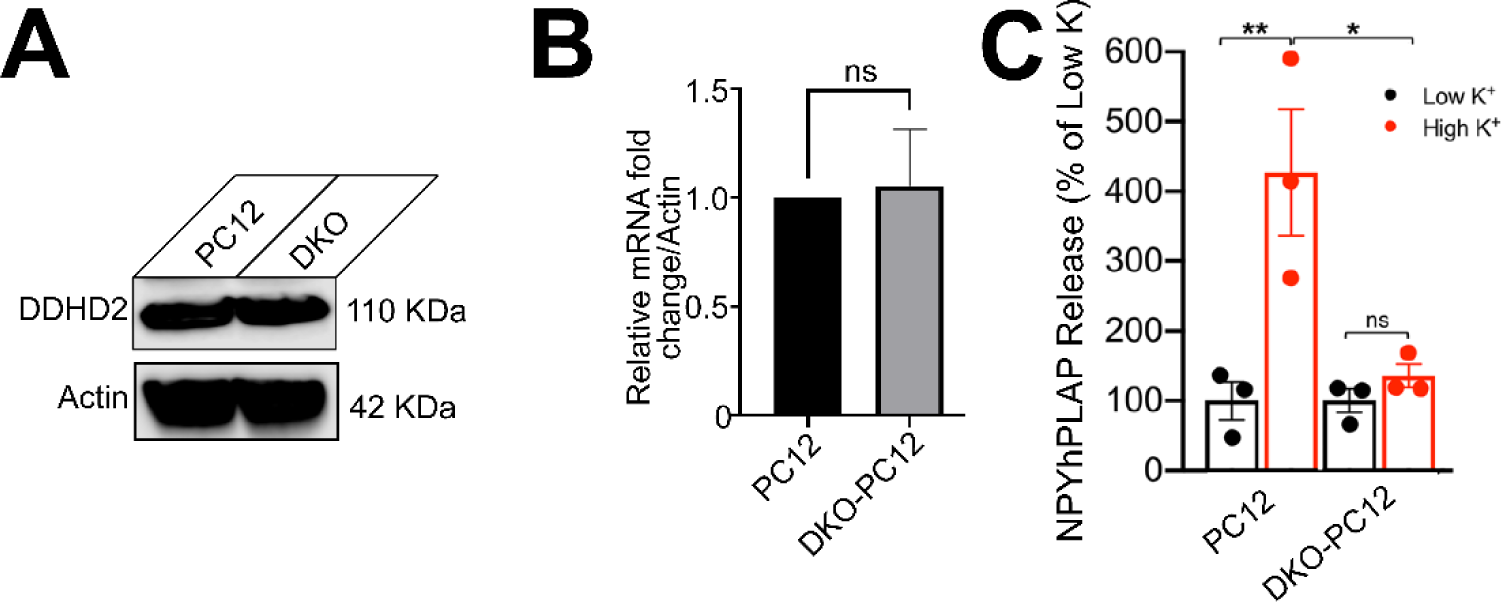
STXBP1 controls the activity-dependent response to exocytosis. **A** Western blot and **B** RT-PCR showing levels of DDHD2 in PC12 and STXBP1/2 DKO cells. Results are shown as mean ± SEM from four independent experiments. **C** Quantification of neuropeptide-Y (NPY) release in PC12 and STXBP1/2 DKO PC12 cells co-transfected with NPY-human placental alkaline phosphatase (NPY-hPLAP) plasmid, and the indicated STXBP1 plasmid, for 72 h. All data are represented as mean ± SEM from 3 independent experiments. One-way ANOVA with Tukey’s correction for multiple comparisons, *p<0.05, **p<0.01, ns = not significant.

**Extended Data Figure 7.**
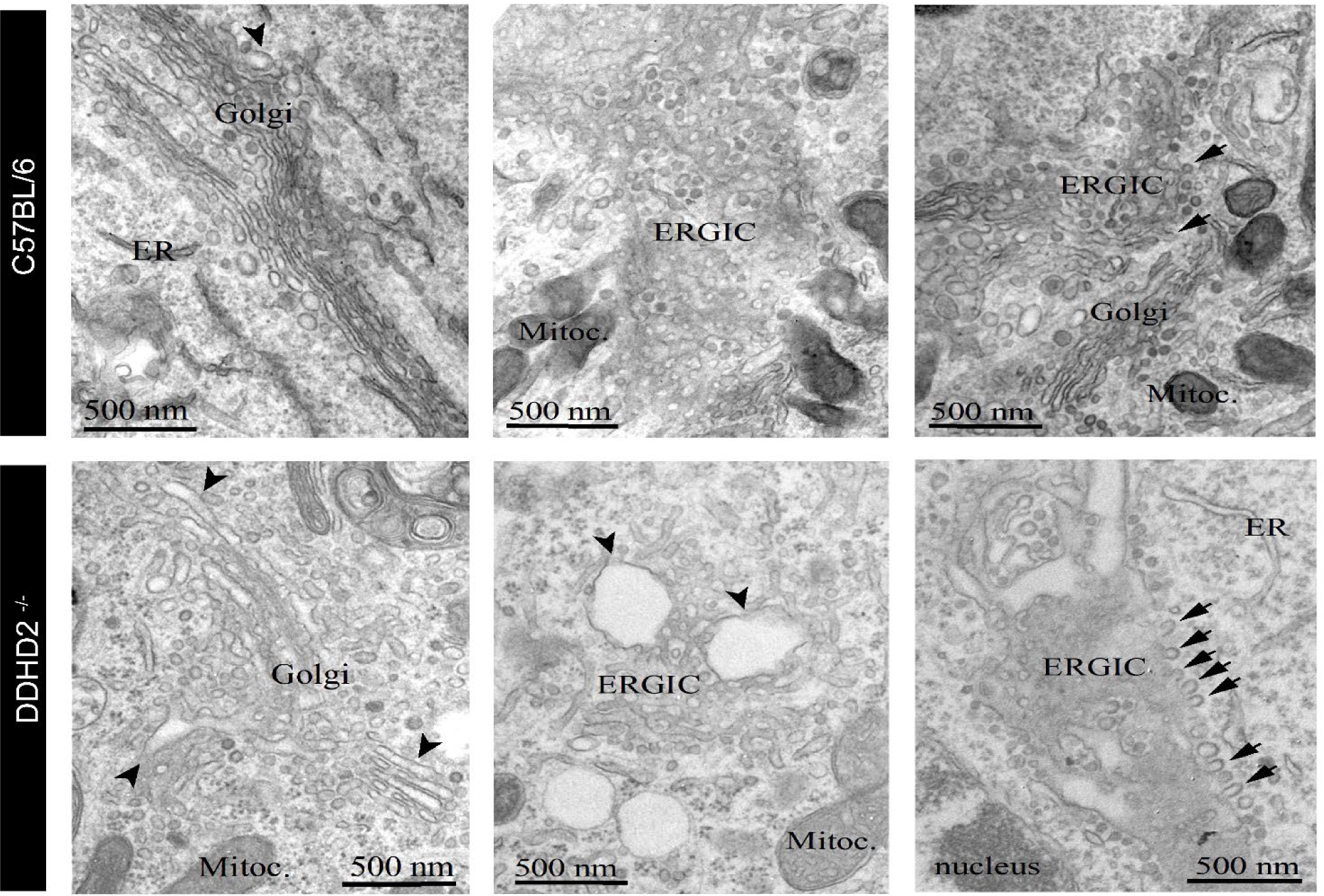
Representative images of the early secretory pathway in *DDHD2^+/+^* vs *DDHD2^-/-^* neurons. Electron micrograph shows dilated lumen of Golgi stacks, enlargement of the ERGIC, with dilated luminal space in the tubulo-vesicular organelle, and arrested vesicles (indicated by arrows) in *DDHD2^-/-^* neurons compared to *DDHD2^+/+^* neurons.

**Extended Data Figure 8.**
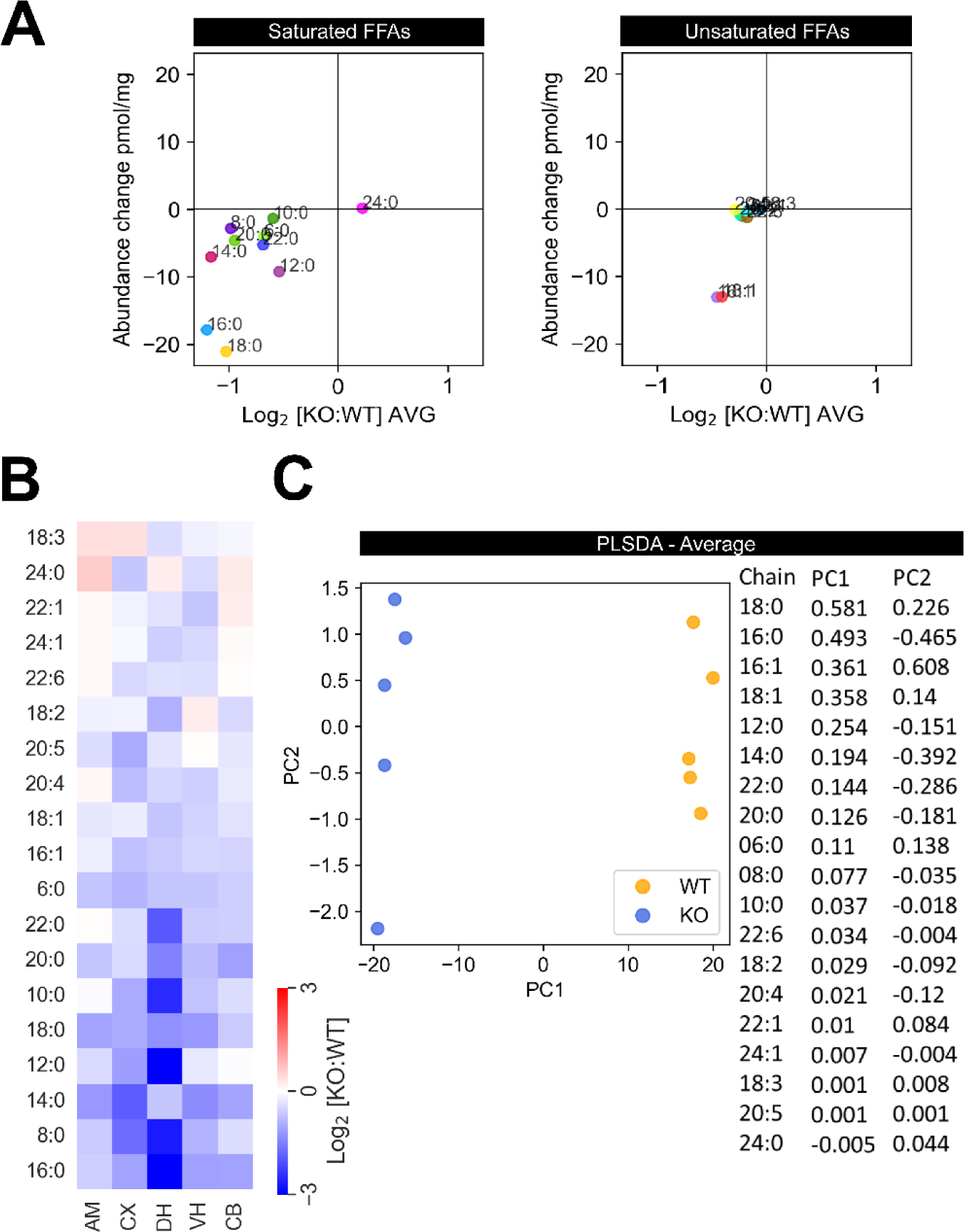
FFA profile of STXBP1 heterozygote mutant mice A. Scatter plot showing absolute vs fold change of average saturated and unsaturated FFAs across the brain of WT versus heterozygote mutant mice. **B** Heatmap of the response of each FFA species in WT versus haploinsufficient STXBP1 animals. **C** Partial least square discriminant analysis (PLSDA) of the average FFA profile across the brain, with tabulated weighting of analyte contribution to clustering shown on the right.

## Extended Data Tables

**Table S1, Related to Figure 1.**
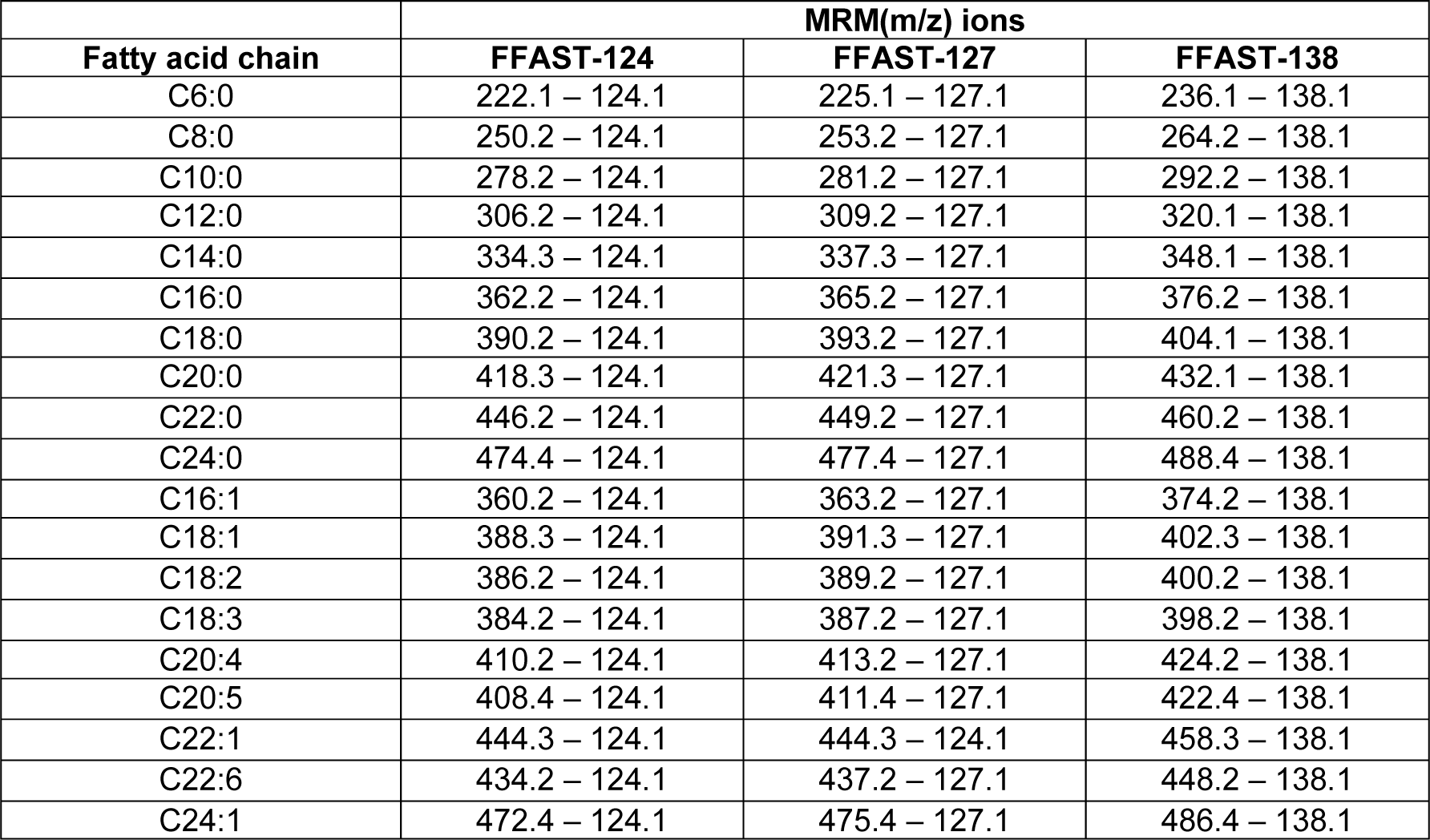
Multiple reaction monitoring (MRM) transitions for the liquid chromatography tandem mass spectrometry (LC-MS/MS) of FFAST derivatized free fatty acids.

**Table S2 Related to Figure 1.**
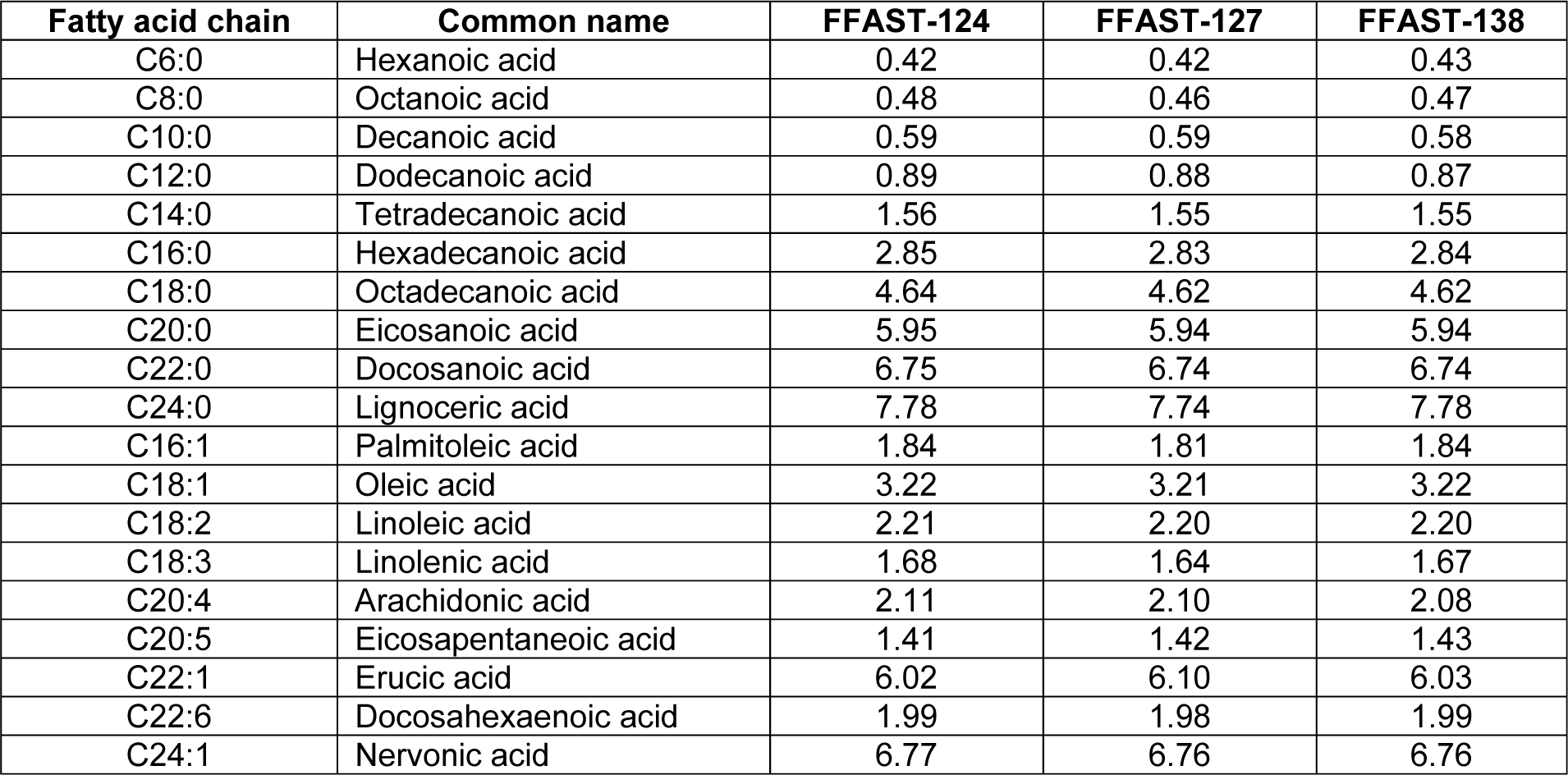
Chromatographic retention times (mins) of FFAST derivatized free fatty acids.

**Table S3 Related to Figure 1.** Multiple Reaction Monitoring (MRM)transition and chromatographic retention times (RT) of FFAST reagents.

| **FFAST Reagents** | **MRM (m/z) ions** | **RT (min)** |
| --- | --- | --- |
| FFAST-124 | 124.1 – 106.2<br>124.1 – 107.1 | 0.24 |
| FFAST-127 | 127.1 – 110.2<br>127.1 – 109.2 | 0.21 |
| FFAST-138 | 138.1 – 106.3<br>138.1 – 121.2 | 0.22 |

**Table S4:**
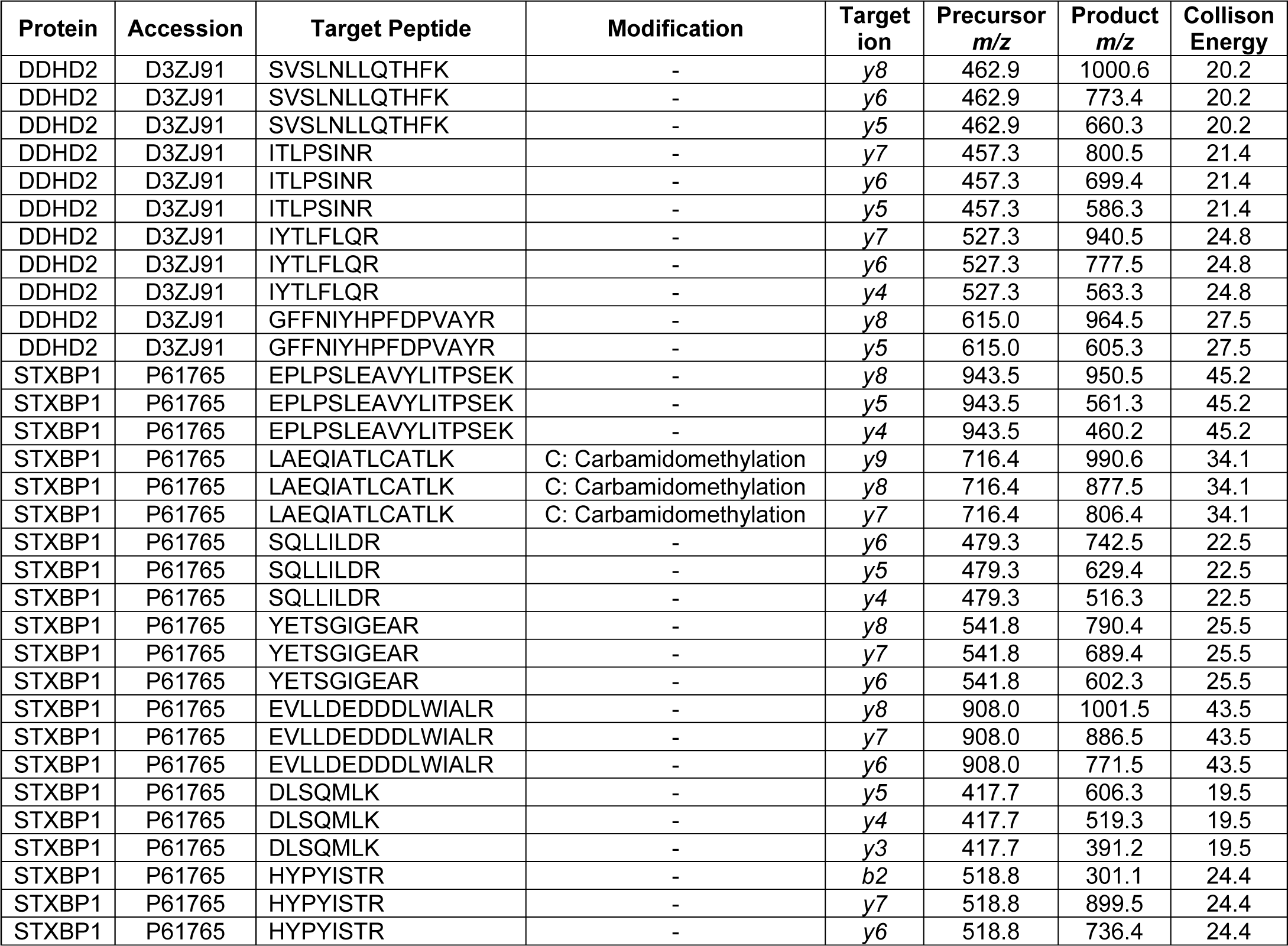
Multiple Reaction Monitoring (MRM) transitions used for the specific quantification of DDHD2 and STXBP1.

**Table S5:** List of proteins detected in the unstimulated PC12 DDHD2 pulldowns.

| Uniprot Accession | Protein Name | Percent Coverage (95% Conf) | Number of Peptides Identified (95% Conf) |
| --- | --- | --- | --- |
| D3ZJ91 | DDHD domain-containing 2 | 80.4 | 206 |
| P61765 | Syntaxin-binding protein 1 | 76.4 | 94 |
| Q6LED0 | Histone H3.1 | 65.4 | 15 |
| D3ZJ08 | Histone H3 | 65.4 | 24 |
| P37805 | Transgelin-3 | 51.3 | 8 |
| P84245 | Histone H3.3 | 55.2 | 13 |
| P11951 | Cytochrome c oxidase subunit 6C-2 | 61.8 | 5 |
| P62630 | Elongation factor 1-alpha 1 | 56.7 | 19 |
| P02262 | Histone H2A type 1 | 57.7 | 11 |
| M0RCL5 | Histone H2A | 57.7 | 8 |
| P62986 | Ubiquitin-60S ribosomal protein L40 | 39.1 | 6 |
| D4AEC0 | Histone H2A | 53.9 | 6 |
| D3ZXP3 | Histone H2A | 52.5 | 12 |
| F1LR10 | LIM domain and actin-binding protein 1 | 39.9 | 20 |
| P49242 | 40S ribosomal protein S3a | 37.1 | 10 |
| M0RBQ5 | Histone H2B | 41.3 | 21 |
| Q4KLJ0 | High mobility group nucleosomal binding domain 2 | 30.0 | 5 |
| P18437 | Non-histone chromosomal protein HMG-17 | 36.7 | 6 |
| A0A0G2JXW4 | Heterogeneous nuclear ribonucleoprotein C | 38.7 | 11 |
| P62982 | Ubiquitin-40S ribosomal protein S27a | 37.8 | 6 |
| P22288 | GTP cyclohydrolase 1 | 35.7 | 6 |
| P85845 | Fascin | 44.2 | 18 |
| F7F9U6 | Plectin | 38.0 | 136 |
| Q78P75 | Dynein light chain 2, cytoplasmic | 44.9 | 3 |
| Q4QQV0 | Tubulin beta chain | 29.8 | 13 |
| D4A9L2 | Serine/arginine-rich splicing factor 1 | 33.9 | 10 |
| P17136 | Small nuclear ribonucleoprotein-associated protein B | 27.7 | 7 |
| A0A0G2K7B3 | Heterogeneous nuclear ribonucleoproteins C1/C2-like | 36.0 | 11 |
| F1LQ48 | Heterogeneous nuclear ribonucleoprotein L | 41.1 | 17 |
| Q5U1W8 | High-mobility group nucleosome binding domain 1 | 43.8 | 10 |
| Q498U4 | SAP domain-containing ribonucleoprotein | 31.9 | 6 |
| A0A0G2K654 | H1.2 linker histone, cluster member | 40.6 | 30 |
| A0A1W2Q680 | H/ACA ribonucleoprotein complex subunit (Fragment) | 40.9 | 6 |
| Q0ZFS8 | RCG61099, isoform CRA_c | 30.5 | 5 |
| Q5RJR8 | Leucine-rich repeat-containing protein 59 | 34.9 | 8 |
| P12839 | Neurofilament medium polypeptide | 31.6 | 24 |
| B0BNK1 | RAB5C, member RAS oncogene family | 22.2 | 4 |
| P55770 | NHP2-like protein 1 | 31.3 | 3 |
| B2RZD1 | Protein transport protein Sec61 subunit beta | 37.5 | 3 |
| P62138 | Serine/threonine-protein phosphatase PP1-alpha catalytic subunit | 33.9 | 10 |
| Q09073 | ADP/ATP translocase 2 | 28.5 | 8 |
| Q9JMJ4 | Pre-mRNA-processing factor 19 | 33.7 | 10 |
| Q5XIF6 | Tubulin alpha-4A chain | 35.7 | 15 |
| P63088 | Serine/threonine-protein phosphatase PP1-gamma catalytic subunit | 34.7 | 9 |
| B0K020 | CDGSH iron-sulfur domain-containing protein 1 | 34.3 | 3 |
| P15865 | Histone H1.4 | 34.3 | 28 |
| M0R4B6 | 60S ribosomal protein L12 | 33.3 | 5 |
| G3V6S0 | Spectrin beta chain | 27.8 | 51 |
| P62142 | Serine/threonine-protein phosphatase PP1-beta catalytic subunit | 32.7 | 9 |
| G3V7U4 | Lamin-B1 | 23.8 | 11 |
| Q91Y94 | Cytochrome c oxidase subunit 4 isoform 2, mitochondrial | 23.3 | 3 |
| A0A140TAB4 | Core histone macro-H2A | 29.1 | 9 |
| D3ZN59 | Similar to High mobility group protein 2 (HMG-2) | 27.3 | 5 |
| Q62733 | Lamina-associated polypeptide 2, isoform beta | 28.1 | 9 |
| F1M3D3 | Heterogeneous nuclear ribonucleoprotein M | 24.7 | 14 |
| Q5PQR0 | RALY heterogeneous nuclear ribonucleoprotein | 24.1 | 8 |
| B5DF91 | ELAV-like protein 1 | 21.2 | 6 |
| P13832 | Myosin regulatory light chain RLC-A | 29.1 | 4 |
| Q5FVM4 | Non-POU domain-containing octamer-binding protein | 21.9 | 7 |
| Q5RJK5 | Chromobox 3 | 29.0 | 3 |
| Q6IFV1 | Keratin, type I cytoskeletal 14 | 17.7 | 10 |
| P43278 | Histone H1.0 | 27.8 | 7 |
| Q6IFU8 | Keratin, type I cytoskeletal 17 | 23.8 | 14 |
| P11240 | Cytochrome c oxidase subunit 5A, mitochondrial | 21.9 | 3 |
| Q6P736 | Polypyrimidine tract-binding protein 1 | 21.4 | 7 |
| P60123 | RuvB-like 1 | 17.1 | 7 |
| Q6PDU1 | Serine/arginine-rich splicing factor 2 | 21.7 | 4 |
| A0A0G2K7X3 | Nuclear ubiquitous casein and cyclin-dependent kinase substrate 1 | 25.5 | 5 |
| D4A720 | RCG61762, isoform CRA_d | 21.4 | 5 |
| A0A0G2K8K0 | Splicing factor proline and glutamine-rich | 19.2 | 10 |
| P62824 | Ras-related protein Rab-3C | 25.1 | 5 |
| Q4KLK7 | NOP56 ribonucleoprotein | 11.4 | 5 |
| P84586 | RNA-binding motif protein, X chromosome retrogene-like | 21.9 | 8 |
| D4A8D5 | Filamin B | 16.0 | 26 |
| F1SW39 | PC4 and SFRS1 interacting protein 1 | 17.4 | 8 |
| Q6P747 | Heterochromatin protein 1-binding protein 3 | 17.7 | 8 |
| Q8K585 | High mobility group protein HMG-I/HMG-Y | 23.4 | 4 |
| Q6AY09 | Heterogeneous nuclear ribonucleoprotein H2 | 16.5 | 5 |
| B2RYU7 | Cbx5 protein | 23.0 | 4 |
| M0R9Z5 | Interferon regulatory factor 2-binding protein 2 | 15.9 | 5 |
| P41777 | Nucleolar and coiled-body phosphoprotein 1 | 19.0 | 12 |
| C0JPT7 | Filamin A | 19.4 | 34 |
| F1M3H8 | Hypothetical protein LOC681410 | 19.3 | 4 |
| A0A0H2UHP9 | RCG39700, isoform CRA_d | 17.8 | 3 |
| M0R9Q1 | RCG48334, isoform CRA_e | 18.2 | 8 |
| D4A9D6 | DEAH box protein 9 | 14.7 | 12 |
| F1LT35 | Similar to 60S ribosomal protein L23a | 19.9 | 3 |
| Q6AY30 | Saccharopine dehydrogenase-like oxidoreductase | 14.0 | 3 |
| P22509 | rRNA 2'- | 17.1 | 4 |
| G3V8V1 | Granulin, isoform CRA_c | 17.8 | 7 |
| P63036 | DnaJ homolog subfamily A member 1 | 18.4 | 4 |
| Q4KM74 | Vesicle-trafficking protein SEC22b | 14.9 | 3 |
| F2Z3T9 | Splicing factor U2AF subunit | 18.5 | 5 |
| A0A0G2JSR7 | Matrin-3 | 15.9 | 10 |
| Q9QZ86 | Nucleolar protein 58 | 15.2 | 6 |
| M0R3M3 | 60S ribosomal protein L7a | 13.7 | 4 |
| D3ZE63 | Uncharacterized protein | 17.4 | 3 |
| A0A0H2UHG4 | Cyclin-dependent kinase 18 | 11.2 | 4 |
| Q5RKJ9 | RAB10, member RAS oncogene family | 16.5 | 3 |
| P19511 | ATP synthase F(0) complex subunit B1, mitochondrial | 13.3 | 3 |
| D4A9A3 | Centromere protein V | 16.4 | 3 |
| M0RCH8 | Ribosomal L1 domain-containing protein 1-like | 7.3 | 3 |
| A0A0G2JSM7 | Adducin 1 (Alpha), isoform CRA_b | 11.0 | 6 |
| A0A0G2JXH6 | Keratin, type II cytoskeletal 73 | 11.5 | 7 |
| D4A250 | Similar to retinoblastoma-binding protein 4 | 8.5 | 4 |
| P81155 | Voltage-dependent anion-selective channel protein 2 | 14.9 | 3 |
| A0A0G2JZR4 | Ras-related protein Rab-11B | 14.7 | 3 |
| D4A0Y6 | Marker of proliferation Ki-67 | 11.5 | 6 |
| Q6AYK1 | RNA-binding protein with serine-rich domain 1 | 14.4 | 4 |
| D3ZCH7 | Adducin 3 (Gamma), isoform CRA_b | 6.8 | 4 |
| B1WC28 | Core histone macro-H2A | 14.2 | 4 |
| Q794E4 | Heterogeneous nuclear ribonucleoprotein F | 10.1 | 3 |
| M3ZCQ2 | U5 small nuclear ribonucleoprotein 200 kDa helicase | 9.6 | 16 |
| G3V741 | Phosphate carrier protein, mitochondrial | 8.4 | 3 |
| Q6AXS3 | Protein DEK | 9.0 | 3 |
| F2Z3Q8 | Importin subunit beta-1 | 10.1 | 6 |
| M0R735 | Heterogeneous nuclear ribonucleoprotein Q | 5.9 | 3 |
| Q91V33 | KH domain-containing, RNA-binding, signal transduction-associated protein 1 | 12.0 | 4 |
| G3V829 | Far upstream element-binding protein 3 | 6.5 | 3 |
| A0A0G2JVB0 | Solute carrier family 2, facilitated glucose transporter member 3 | 8.9 | 3 |
| Q1LZ53 | DNA (cytosine-5)-methyltransferase 3A | 6.9 | 6 |
| D4ACS0 | Protein phosphatase 1 regulatory subunit | 8.2 | 6 |
| A0A0H2UHL9 | Drebrin | 8.6 | 5 |
| D4A206 | Treacle ribosome biogenesis factor 1 | 9.3 | 9 |
| G3V769 | E3 ubiquitin-protein ligase TRIM35 | 8.7 | 3 |
| P85834 | Elongation factor Tu, mitochondrial | 10.0 | 3 |
| Q04931 | FACT complex subunit SSRP1 | 7.6 | 5 |
| Q8R4T2 | E3 ubiquitin-protein ligase SIAH2 | 8.0 | 3 |
| G3V624 | Coronin | 7.6 | 3 |
| D4A4J0 | FACT complex subunit | 7.3 | 5 |
| Q566E4 | Heterogeneous nuclear ribonucleoprotein R | 5.2 | 3 |
| O35274 | Neurabin-2 | 7.1 | 4 |
| G3V9Y1 | Myosin, heavy polypeptide 10, non-muscle, isoform CRA_b | 5.0 | 7 |
| A0A0G2K2M9 | Serine/arginine repetitive matrix 2 | 6.5 | 13 |
| O35821 | Myb-binding protein 1A | 5.8 | 5 |
| F7FF45 | Nuclear mitotic apparatus protein 1 | 5.0 | 8 |
| A0A0G2K5U9 | Alpha-actinin-4 | 5.3 | 4 |
| G3V661 | Bromodomain adjacent to zinc finger domain protein 1B | 2.7 | 4 |
| F1M4A0 | Tight junction protein 1 (Predicted) | 6.2 | 8 |
| M0R965 | RNA helicase | 5.5 | 4 |
| F1LQZ9 | Microtubule-associated protein 6 | 6.4 | 3 |
| M0RBF0 | Scaffold attachment factor B1 | 5.3 | 3 |
| F1LNL2 | SWI/SNF related, matrix associated, actin dependent regulator of chromatin, subfamily a, member 5 (Predicted), isoform CRA_a | 3.3 | 3 |
| A0A0G2JWP8 | DNA topoisomerase 2 | 6.0 | 8 |
| P27008 | Poly [ADP-ribose] polymerase 1 | 3.5 | 3 |
| E9PT66 | Splicing factor 3b, subunit 3 | 3.0 | 3 |
| E9PU01 | DNA helicase | 1.4 | 3 |
| M0R9L0 | Nascent polypeptide-associated complex subunit alpha | 1.9 | 3 |
| E9PST5 | Apoptotic chromatin condensation inducer 1 | 2.6 | 3 |

